# Nonlinear control of transcription through enhancer-promoter interactions

**DOI:** 10.1101/2021.04.22.440891

**Authors:** Jessica Zuin, Gregory Roth, Yinxiu Zhan, Julie Cramard, Josef Redolfi, Ewa Piskadlo, Pia Mach, Mariya Kryzhanovska, Gergely Tihanyi, Hubertus Kohler, Peter Meister, Sebastien Smallwood, Luca Giorgetti

## Abstract

Chromosome structure in mammals is thought to regulate transcription by modulating the three-dimensional interactions between enhancers and promoters, notably through CTCF-mediated interactions and topologically associating domains (TADs)^1–4^. However, how chromosome interactions are actually translated into transcriptional outputs remains unclear. To address this question we use a novel assay to position an enhancer at a large number of densely spaced chromosomal locations relative to a fixed promoter, and measure promoter output and interactions within a genomic region with minimal regulatory and structural complexity. Quantitative analysis of hundreds of cell lines reveal that the transcriptional effect of an enhancer depends on its contact probabilities with the promoter through a non-linear relationship. Mathematical modeling and validation against experimental data further provide evidence that nonlinearity arises from transient enhancer-promoter interactions being memorized into longer-lived promoter states in individual cells, thus uncoupling the temporal dynamics of interactions from those of transcription. This uncovers a potential mechanism for how enhancers control transcription across large genomic distances despite rarely meeting their target promoters, and for how TAD boundaries can block distal enhancers. We finally show that enhancer strength additionally determines not only absolute transcription levels, but also the sensitivity of a promoter to CTCF-mediated functional insulation. Our unbiased, systematic and quantitative measurements establish general principles for the context-dependent role of chromosome structure in long-range transcriptional regulation.

## Main text

Transcriptional control critically depends on distal *cis*-regulatory elements such as enhancers, which control the tissue specificity and developmental timing of a large number of genes^5^. Enhancers are often located hundreds of kilobases away from their target promoters and are thought to control gene expression by interacting with promoters in the three-dimensional space of the nucleus. However, the biochemical mechanisms leading to the exchange of regulatory information as well as the radius at which it occurs remain poorly understood^6,7^. Chromosome conformation capture (3C) methods such as Hi-C, which measure physical proximity between genomic sequences using chemical crosslinking and ligation^8^, have shown that interactions between enhancers and promoters predominantly occur within sub-megabase domains known as topologically associating domains (TADs). TADs mainly arise from nested interactions between sites that are bound by the DNA-binding protein CTCF which act as barriers for the loop extrusion activity of the cohesin complex^9^.

TAD boundaries and CTCF loops are thought to contribute to the regulation of gene expression by favoring enhancer-promoter communication within subsets of genomic regions and disfavoring it with respect to surrounding genomic sequences. Topological constraints have indeed been shown to be able to functionally insulate regulatory sequences^1,3,4,10^. Recently, however, this view has been challenged by reports showing that disruption of TAD boundaries^11,12^ or global depletion of CTCF and cohesin^13,14^ do not lead to systematic large-scale changes in gene expression, and that some regulatory sequences can act across TAD boundaries^15^. Inversions, deletions, and insertions of single CTCF sites also have been reported to result in variable effects on gene expression, ranging from major^2,4,16–18^ to very moderate or none^12,19,20^. The very notion that physical proximity is required to elicit transcriptional regulation has been questioned by the observed lack of correlation between transcription and proximity in single cells^21,22^. As a consequence of these variable and locus-dependent results, it is highly debated if there are indeed general principles that determine how physical interactions enable enhancer action^23^; and under which conditions TAD boundaries and CTCF sites manage to insulate regulatory sequences. Enhancer-promoter genomic distance might additionally contribute to transcriptional regulation^24,25^. It is however unclear if the efficiency of promoter regulation by an enhancer is uniform within a single TAD^26,27^, or if transcription rather depends on enhancer-promoter genomic distance within TAD boundaries^25,28^.

Addressing these fundamental questions requires a quantitative understanding of the relationship between transcriptional output and enhancer-promoter interactions, in conditions where confounding effects by additional regulatory and structural interactions can be minimized. Here we provide such a description using a novel experimental approach that enables systematic measurement of promoter activity as a function of its genomic distance with an enhancer. Coupled with a mathematical model of enhancer-promoter communication this reveals that interactions with an enhancer determine promoter output in a non-linear manner, and that transcription levels as well as functional insulation by topological constraints depend on enhancer strength and distance from a promoter.

To systematically test enhancer function we established an assay in which an enhancer can be mobilized from an initial genomic location and reinserted at a large number of different genomic locations with respect to its cognate promoter. This enables the measurement of transcription levels as a function of enhancer location, and hence of enhancer-promoter contact frequencies (**Fig. 1A**). More specifically we generated mouse embryonic stem cells (mESCs) carrying a transgene where a promoter drives the expression of enhanced green fluorescent protein (eGFP). The eGFP transcript is split in two by a piggyBac transposon that contains the promoter’s cognate enhancer (**Fig. 1B**) and prevents translation of a functional eGFP protein. Upon expression of the PBase transposase, the transposon is excised and reintegrated randomly in the genome but preferentially in vicinity of the site of excision^29^. The excision leads to the reconstitution of a functional eGFP transcript whose expression levels are used to isolate clonal cell lines by sorting single eGFP positive (eGFP+) cells (**Fig. 1C-D**). This allows the rapid generation of hundreds of cell lines, in each of which the enhancer exerts its regulatory activity from a single distinct genomic position. The enhancer genomic position and eGFP expression levels in every cell line are then determined enabling the measurement of transcription level as a function of enhancer location (**Fig. 1D**).

**Figure 1.**
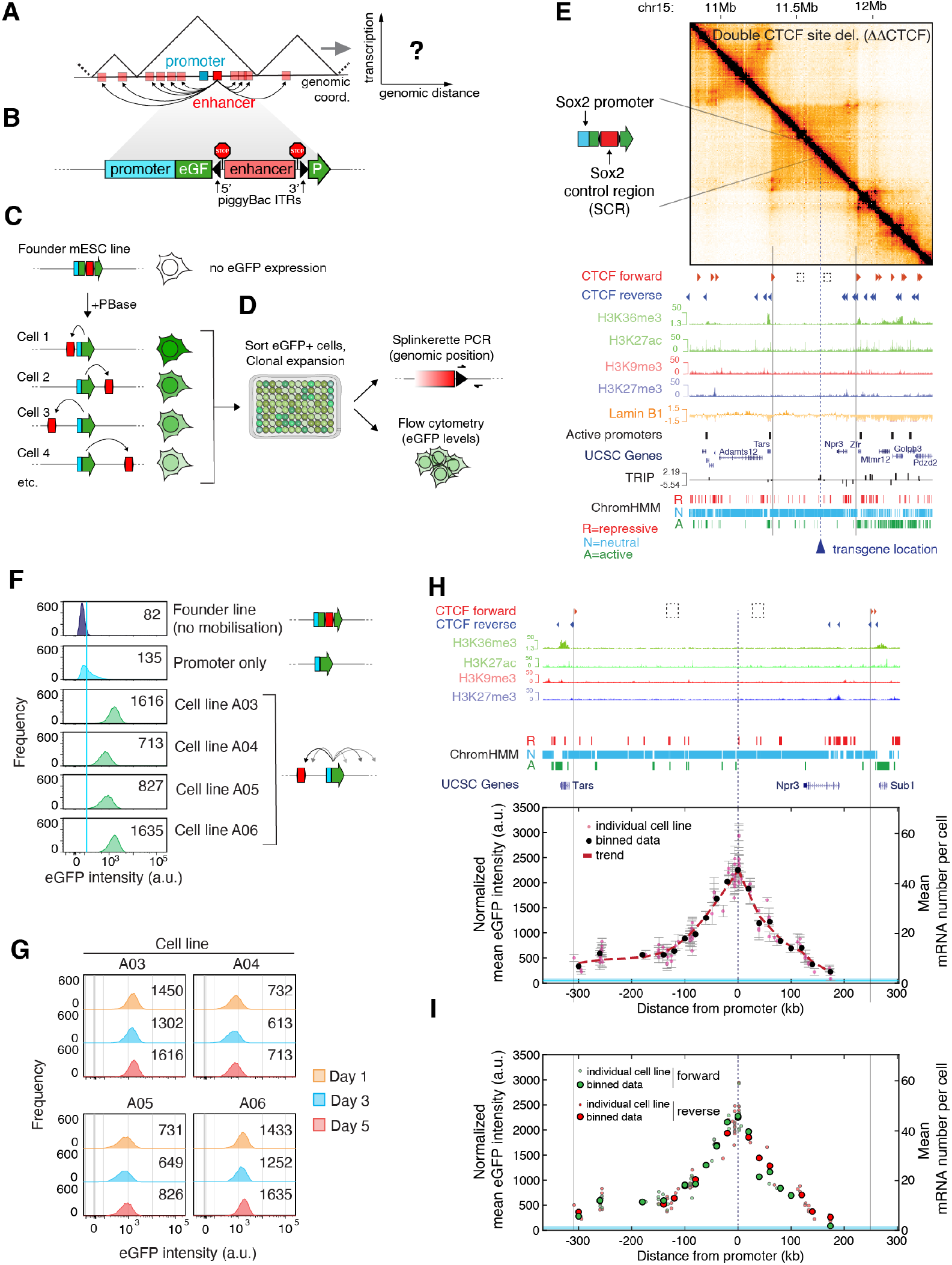
Enhancer action is modulated by genomic distance from the target promoter and constrained by TAD boundaries. A. Scheme of the experimental strategy. Mobilisation of an enhancer around its target promoter allows measuring transcription as a function of genomic distance to the enhancer. B. Structure of the transgene developed for this study. A mammalian promoter drives transcription of a split eGFP gene containing a piggyBac transposon cassette. The transposon harbors an enhancer flanked on each side by a stop cassette composed of an SV40 poly-adenylation site and a transcriptional pause signal from the human alpha2 globin gene to minimize potential transcription originating inside the enhancer. The PBase transposase scarlessly excises sequences between the two piggyBac inverted terminal repeats (ITRs), including the ITRs. C. The piggyBac-enhancer cassette is excised upon expression of PBase transposase and reinserted at random genomic locations. Insertions that lead to promoter activation result in eGFP levels distinct from the background detected in cells where the transposon is not excised. D. Fluorescence-activated cell sorting (FACS) and clonal expansion of single eGFP+ cells allow the rapid generation of large numbers of cell lines where the enhancer is inserted at a single genomic position and drives promoter transcription. Splinkerette PCR and flow cytometry are then used to determine the enhancer genomic position and expression levels of the promoter. E. Capture Hi-C contact map at 6.4-kb resolution following the deletion of both internal CTCF motifs, plotted with genomic datasets in mESCs in a 2.6-Mb region centered around the TAD we used for the experiments, highlighting simple chromosome structure and low baseline genomic activity within the TAD. Vertical grey lines: TAD boundaries. Dashed blue line: genomic position of the integrated transgene carrying the *Sox2* promoter and enhancer (*Sox2* control region, SCR). Dashed squares indicate the position of the two deleted CTCF sites (cf. Suppl. Fig. S1B-C). F. Representative flow cytometry profiles from the founder mESC line, a control cell line where eGFP transcription is driven from the same genomic location by the *Sox2* promoter alone, and four eGFP+ cell lines where the SCR was mobilised and reinserted. Solid light blue line: mean eGFP levels in promoter-only line. Numbers indicate median eGFP values. G. eGFP levels in individual eGFP+ cell lines over cell passages. Numbers indicate median eGFP values. H. Normalized mean eGFP intensities in individual eGFP+ cell lines are plotted as a function of the genomic position of the SCR. Data from 112 individual cell lines (light red dots) from a single experiment (error bars: standard deviation of three measurements performed on different days, as in panel G) and average eGFP values calculated within equally spaced 20-kb bins (black dots) are shown. Dashed red line: trendline based on a smoothing spline interpolant of the average eGFP values. Mean mRNA numbers per cell were inferred from eGFP counts using calibration with smRNA FISH, see Suppl. Fig. 1H. Shaded light blue area indicates the interval between mean +/- standard deviation of eGFP levels in three promoter-only cell lines. I. Same data as in panel H. Single clone eGFP levels and binned data are colored according to the genomic orientation of the SCR.

To minimize confounding effects from additional regulatory sequences and structural interactions, we integrated the transgene within a 560 kb TAD located on chromosome 15 which carries minimal regulatory and structural complexity. This TAD does not contain expressed genes or active enhancers, is mostly composed of ‘neutral’ chromatin states (identified using chromHMM^30^) except for a repressive ~80 kb region at its 3’ side (**Suppl. Fig. 1A**) and displays minimal sub-TAD structure mediated by two internal forward CTCF sites (**Suppl. Fig. 1A-B**). To further decrease the TAD’s structural complexity, we homozygously deleted the two internal CTCF sites which led to the loss of the associated loops observed by Capture Hi-C (c-HiC) (**Suppl. Fig. 1C**) and resulted in a TAD with a simple homogeneous internal structure (**Fig. 1E** and **Suppl. Fig.1C**).

We first integrated a version of the transgene carrying the mouse *Sox2* promoter and the essential 4.8-kb region of its distal enhancer^31^ known as *Sox2* control region (SCR)^31,32^ (**Supplementary Fig. 1D, Methods**), from which we deleted its single CTCF site which is not essential for transcriptional regulation at the endogenous locus^19^. Using targeted nanopore sequencing with Cas9-guided adapter ligation (nCATS)^33^, we verified that the transgene was inserted intact and in a single copy (**Suppl. Fig. 1E**). Transgene insertion did not lead to significant structural rearrangements within the TAD besides the formation of new moderate interactions between the transgene and the CTCF sites at the 3’ and 5’ end of the TAD (**Suppl. Fig. 1F**).

Mobilization of the piggyBac-SCR cassette led to random genomic reinsertion events with a preference for chromosome 15 where it was initially located (**Suppl. Fig. 1G**), as previously observed^29^. Single experiments resulted in several tens of eGFP+ cell lines whose eGFP levels were unimodally distributed (**Figure 1F**), higher than those detected in control lines where transcription was driven by the *Sox2* promoter alone (**Figure 1F**), and remained stable over several cell passages (**Fig. 1G**). Mean eGFP levels in single cell lines were also linearly correlated with the average number of eGFP mRNAs measured using singlemolecule RNA fluorescent in situ hybridisation (smRNA FISH) (**Suppl. Fig. 1H**). We thus used mean flow cytometry eGFP values as a direct readout of promoter transcriptional activity.

Unbiased mapping of piggyBac-SCR positions in more than two hundred eGFP+ cell lines revealed that in 99.6% of them (240/241) the enhancer had reinserted within the TAD where the reporter is located (**Fig. 1H**, replicate experiment in **Suppl. Fig. 1I**). We isolated a single cell line where the enhancer was transposed outside the TAD. In this case, eGFP levels were actually comparable to promoter-only control cells (**Suppl. Fig. 1J**). Strikingly, within the TAD expression levels quantitatively depended on the actual position of the enhancer and monotonically decreased with increasing enhancer-promoter genomic distance (**Fig. 1H**). Varying the genomic distance between the enhancer and the promoter within the domain accounted for a ten-fold dynamic range in gene expression, from approximately 4.5 to 45 mRNAs per cell on average based on calibration with smRNA FISH (**Suppl. Fig. 1H**). Transcription levels decreased symmetrically on both sides of the ectopic promoter, except for insertions downstream of the nontranscribed *Npr3* gene that did not generate detectable transcription levels (**Fig. 1H**) possibly due to the influence of the flanking repressive region. Mild positive and negative deviations from the average decay in transcription levels were indeed correlated with local enrichment in active and repressive chromatin states, respectively, surrounding the SCR insertion sites (**Suppl. Fig. 1K**). In line with the classical notion derived from reporter assays that enhancer activity is independent of its genomic orientation^34^, we additionally found no differences in transcription levels generated by enhancers inserted in forward or reverse orientations (**Fig. 1I**).

These data show that the range of activity of the SCR enhancer is delimited by TAD boundaries. With the exception of the short repressive region at the 3’-end, the entire TAD is permissive for communication between this strong enhancer and the ectopic *Sox2* promoter. Transcription levels however quantitatively depend on enhancer-promoter genomic distance within the domain.

To understand how transcription levels inside the TAD and insulation at its boundaries are generated, we next examined the relationship between transcription levels and contact probabilities. Although reads from the wild-type non-targeted allele might underemphasize structural changes introduced by the heterozygous insertion of the transgene, contact patterns detected in cHi-C did not change markedly in individual clones where the SCR was reinserted in the TAD compared to the founder line before piggyBac mobilisation (**Suppl. Fig. 2A**). This suggests that the ectopic *Sox2* enhancer and promoter do not create prominent specific interactions. To infer contact probabilities between promoter and enhancer locations, we thus normalized cHi-C data from the founder line at 6.4-kb resolution setting the average counts from adjacent genomic bins correspond to a contact probability of one^35^ (**Fig. 2A**).

**Figure 2.**
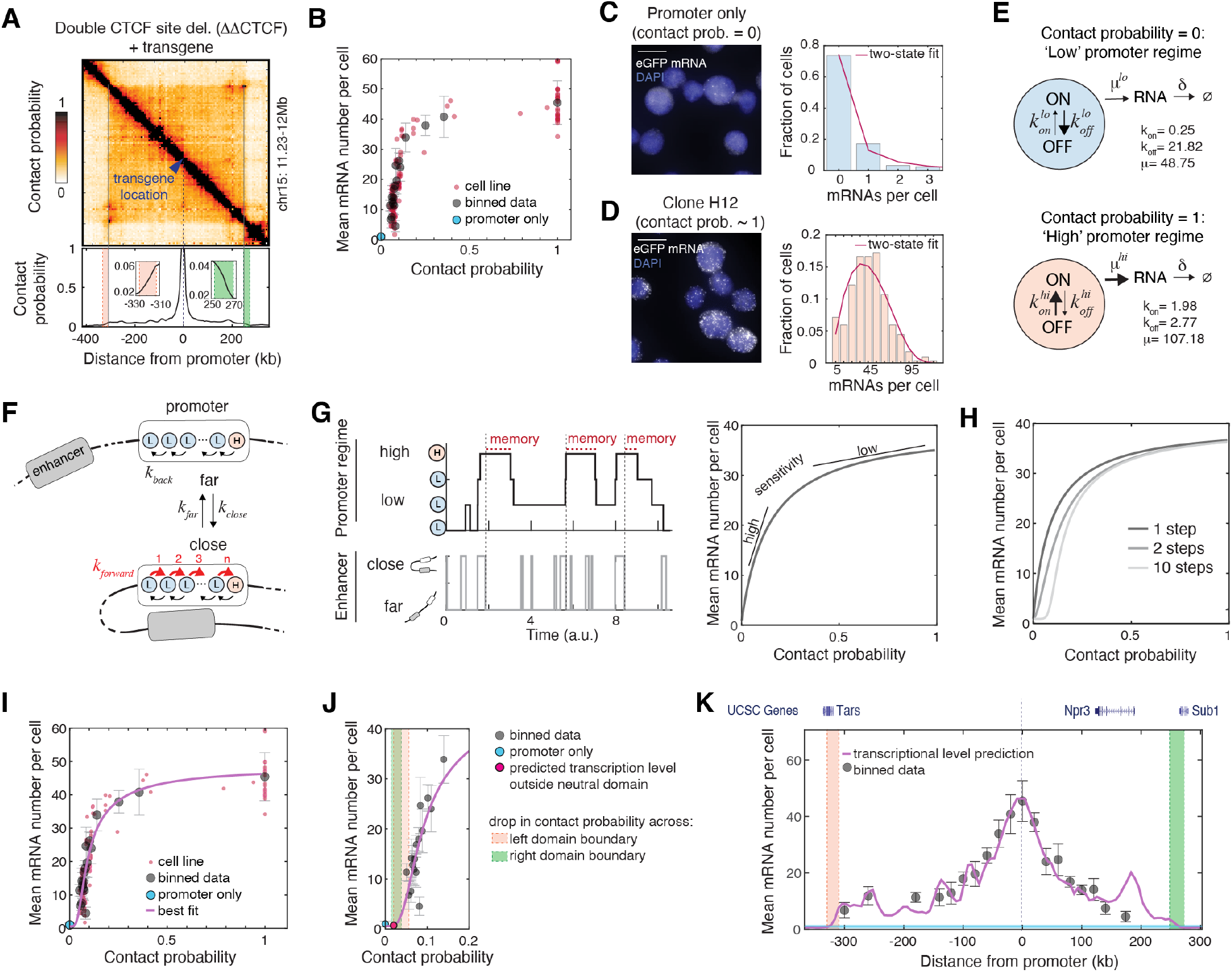
Enhancer-promoter contacts are nonlinearly translated into transcription through a small number of rate-limiting regulatory steps and memorized into long-lived promoter states. A. Top: Capture Hi-C data (6.4-kb resolution) from the founder cell line used for experiments in Figure 1. Read counts were transformed into contact probabilities as described in the main text and Methods section. Bottom: Cross section showing contact probabilities from the location of the ectopic *Sox2* transgene. Insets: zoom-in of the drop in contact probability across TAD boundary regions, detected using CaTCH on Hi-C from Redolfi et al.^37^. B. Inferred mean GFP mRNA numbers per cell are plotted against contact probabilities between the ectopic *Sox2* promoter and the locations of SCR insertions. mRNA numbers were inferred from flow cytometry using smRNA FISH calibration, Suppl. Fig. 1H. Individual cell lines (error bars as in fig. 1H) are plotted together with eGFP average values calculated in equally spaced genomic bins as in Figure 1H. C. Representative smRNA FISH image and distribution of mRNA counts per cell in mESCs where eGFP transcription is driven by the ectopic *Sox2* promoter alone. Line: fit with the two-state promoter model shown in Fig. 2E, top. D. Same as C but in a clonal cell line where the SCR is in the immediate vicinity of the ectopic *Sox2* promoter (−6.1 kb). Line: fit with the two-state promoter model shown in Fig. 2E, bottom. E. Scheme and parameters of the two-state models used to fit cell lines in panels C and D. F. Schematic description of the mathematical model of enhancer-promoter communication. When the enhancer is close to the promoter (bottom), a series of n reversible regulatory steps occur with rates kforward and kback during which the promoter remains in the low (L) two-state regime of Fig. 2E. After the n-th step, the promoter transiently switches to the high (H) two-state regime. When the enhancer is far from the promoter (top), the regulatory steps can only be reversed (with rate kback). G. Left panel: Representative example of single-cell dynamics of enhancer states and promoter regimes predicted by the model. Memory is shown as the time that the promoter stays in the high two-state regime (H) after the last contact is disassembled. Right panel: Representative steady-state transcription levels as a function of enhancer-promoter contact probabilities predicted by a model with long memory and efficient intermediate regulatory steps. H. Representative model prediction for steady-state transcription levels as a function of enhancer-promoter contact probabilities 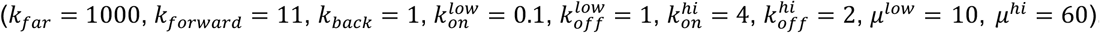. Increasing the number of pre-activation regulatory steps increases the sigmoidality of the transcriptional response. I. Best fit to the experimental data of panel B. Best-fit parameters are shown in Suppl. Fig. 3A. J. Close-up view of panel I highlighting model behavior at low contact probabilities and predicted insulation outside TAD boundaries. Drops in contact probabilities across TAD boundaries are highlighted by red and green shaded areas. K. Model prediction (purple line) of transcription levels generated by the SCR within the TAD, plotted against binned expression data (cf. Fig. 1H). Drops in contact probabilities across TAD boundaries are highlighted by red and green shaded areas as in panel J. Shaded light blue area indicates the interval between mean +/- standard deviation of eGFP levels in three promoter-only cell lines.

We observed that contact probabilities steeply decayed with increasing genomic distance from the promoter, fell drastically while approaching the TAD boundaries (from 1 to 0.06 and 0.04 inside left and right boundaries, respectively), and further dropped by a factor ~3 across boundaries (2.9- and 2.7-fold for left and right boundaries, respectively) (**Fig. 2A**). This is in line with previous estimations^36^ confirmed by crosslinking and ligation-free methods^37^ and is representative of contact probabilities experienced by active promoters genome-wide in mESCs (**Suppl. Fig. 2B-C**). Such trend is however at odds with our observation that transcription levels rather mildly decreased inside the TAD and dropped to promoter-only levels outside its boundaries (**Fig. 1H** and **Suppl. Fig. 2D**). Interestingly, plotting mean eGFP mRNA numbers inferred from flow cytometry calibrated with smRNA FISH as a function of contact probabilities revealed indeed a highly non-linear relationship (**Fig. 2B**). Thus, if physical interactions with an enhancer determine transcriptional output at the promoter, a mechanism must exist that converts their contact probabilities nonlinearly into transcription levels.

We sought to understand if such a nonlinear relationship could be related to how enhancer-promoter interactions translate into transcriptional events in individual cells. Transcription occurs as the endpoint of multiple kinetically distinct processes at the promoter, which involve small numbers of molecules and result in intermittent transcription bursts^38^. Bursty promoter behavior can be described in terms of a two-state model of gene expression^39^, in which the promoter stochastically switches with rates *k_on_* and *k_off_* between an OFF (inactive) and an ON (active) state where transcription can occur with rate *μ*. In line with this notion, the eGFP mRNA distribution measured by smRNA FISH in control cell lines where the ectopic *Sox2* promoter drives transcription alone (and thus its contact probability with the enhancer is zero by definition) was well approximated by a ‘low-regime’ two-state model with low transcriptional activity (**Fig. 2C, 2E** top panel). In cell lines where the SCR is adjacent to the promoter and their contact probability is close to one, mRNA distributions were rather described by a ‘high-regime’ two-state model with higher transcriptional activity (**Fig. 2D, 2E** bottom panel). The observed nonlinear relationship thus might represent a gradual and nonlinear conversion of promoter operation from a ‘low’ to a ‘high’ two-state regime as contact probabilities with its enhancer are increased from zero to one (**Fig. 2E**).

We next asked which mechanisms would account for such nonlinear conversion. A simple model where each transcription burst is the result of a single promoter-enhancer interaction would generate a linear conversion of contact probabilities into transcription levels and thus would fail to account for the nonlinear behavior we observed. We however reasoned that the molecular processes mediating enhancer-promoter communication (e.g. recruitment of transcription factors and coactivators, assembly of the Mediator complex^6^) are also likely to introduce rate-limiting steps. Coupled to the intrinsic stochastic dynamics of the promoter, those steps could in principle generate nonlinear conversion of contact probabilities into transcription levels. To explore this concept quantitatively we developed a mathematical model of enhancer-promoter communication. Consistent with the eGFP mRNA distribution measured by FISH in cells lacking the SCR (**Fig. 2C**), we assumed that in the complete absence of interactions with an enhancer, the promoter operates as a low-regime (L) two-state model. Stochastic interactions with an enhancer occur and disassemble with rates *k_close_* and *k_far_*, respectively, and trigger one or more (*n*) kinetically distinct, reversible regulatory steps that transmit regulatory information to the promoter (**Fig. 2F**). At the end of these intermediate steps, which occur with rates *k_forward_* and *k_back_*, the promoter transiently switches into a high-regime (H) two-state model with modified *on*, *off*, and initiation rates and thus transiently increases its transcriptional activity (**Fig. 2F**). In the limit case where promoter and enhancer are always in contact, the promoter mainly operates in the high two-state regime, in agreement with mRNA distributions measured when the SCR is in the immediate vicinity of the *Sox2* promoter (**Fig. 2D, 2E**).

Strikingly, analytical solution of the model (**Suppl. Model Description**) showed that under these hypotheses contact probabilities are transformed into transcription levels in a non-linear fashion, irrespective of parameter values. Non-linearity arises from the ability of the promoter to remain in the high-regime mode longer than the duration of interactions with the enhancer, thus ‘memorizing’ stochastic interactions (**Fig. 2G**). If memory is long (i.e. the promoter remains in the high-regime mode much longer than the average duration of an interaction, *k_back_* ≪ *k_far_*), and if forward transitions through intermediate regulatory steps are favored over backward reactions (*k_forward_* > *k_back_*), then transcription levels become highly sensitive to changes in contact probabilities when contact probabilities are low, and conversely poorly sensitive when contact probabilities are high (**Fig. 2G**), in qualitative agreement with our experimental observations. Increasing the number *n* of intermediate regulatory steps introduces an increasingly stronger kinetic barrier in the transmission of regulatory information, which makes the transition to the ‘high’ regime mode difficult at low contact probabilities. The transcriptional response becomes thus sigmoidal (**Fig. 2H**). Importantly our model does not require any hypotheses on what actually drives enhancer-promoter interactions, nor on their specific molecular range. Encounters might occur through random collisions of the chromatin fiber or the loop extrusion activity of cohesin, and might either involve molecular-range interactions or be mediated by molecular condensates.

We next fitted the model simultaneously to the mRNA FISH distributions shown in **Fig. 2C-D** and to the experimental transcriptional response of **Fig. 2B** (see **Methods**). This revealed that the best agreement with the data occurred with four to eleven intermediate regulatory steps (**Suppl. Model Description**), corresponding to a sigmoidal curve (**Fig. 2I**, shown with six intermediate steps; best-fit parameters in **Suppl. Fig. 3A**). In these conditions, the model predicts that the SCR is no longer able to activate the *Sox2* promoter once it crosses the ~3-fold drop in contact probabilities generated by the TAD boundaries (**Fig. 2J**) and accurately reproduces transcription levels within the domain (**Fig. 2K**). In contrast, fitting with a model with a single intermediate step resulted in a lower agreement with the experimental data (**Suppl. Fig. 3B-C**), as well as qualitatively worse predictions of transcription levels within and outside TAD boundaries (**Suppl. Fig. 3D**). Best-fit parameters correspond to scenarios where enhancer-promoter interactions are short (~1/20 of mRNA life-time, which is the elementary time unit in the model and should be in the order of 1.5 hours^40^). In these conditions, promoter-enhancer contacts and transcription bursts should be temporally uncorrelated despite being causally linked (**Suppl. Fig. 3E**; correlation < 0.002, **Suppl. Fig. 3F**), as recently observed at the endogenous *Sox2* locus^22^. The model predicts that both promoter burst frequency and burst size depend on contact probabilities (**Suppl. Fig. 3G-I**). Interestingly, however, within the TAD, enhancer-promoter contact probabilities are in a range where mainly burst frequency is affected (**Suppl. Fig. 3J-K**), in line with previous reports that enhancers mainly modulate burst frequency^24,41^.

Analysis of experimental data through our model thus suggests that conversion of enhancer-promoter contact probabilities into transcription levels occurs through a small number of rate-limiting regulatory steps following transient enhancer-promoter interactions, which are then memorized into longer-lived promoter states. The nonlinear response generated by this mechanism readily explains both why sharp decays in contact probabilities at small genomic distances from the promoter are buffered in milder changes in transcription levels, thus allowing an enhancer to act from large genomic distances, and why enhancer action is restricted by TAD boundaries.

Enhancers have been previously suggested to be able to modulate the probability that a gene is transcribed in individual cells, rather than its absolute level of expression^42^. We thus investigated whether enhancer position within the TAD would also affect cell-to-cell transcriptional heterogeneity within individual cell lines. We observed that in eGFP+ cell lines, coefficients of variation of eGFP signals assessed by flow cytometry (CV, standard deviation divided by mean) increased as a function of increasing genomic distances between the SCR and the ectopic *Sox2* promoter (**Fig. 3A**). Cell lines where the enhancer was located hundreds of kb away from the promoter showed broader and asymmetric distributions of eGFP signals compared to cell lines where the SCR is located near the promoter, with different fractions of cells expressing eGFP levels either below or above the mean expression level (**Fig. 3B**).

**Figure 3.**
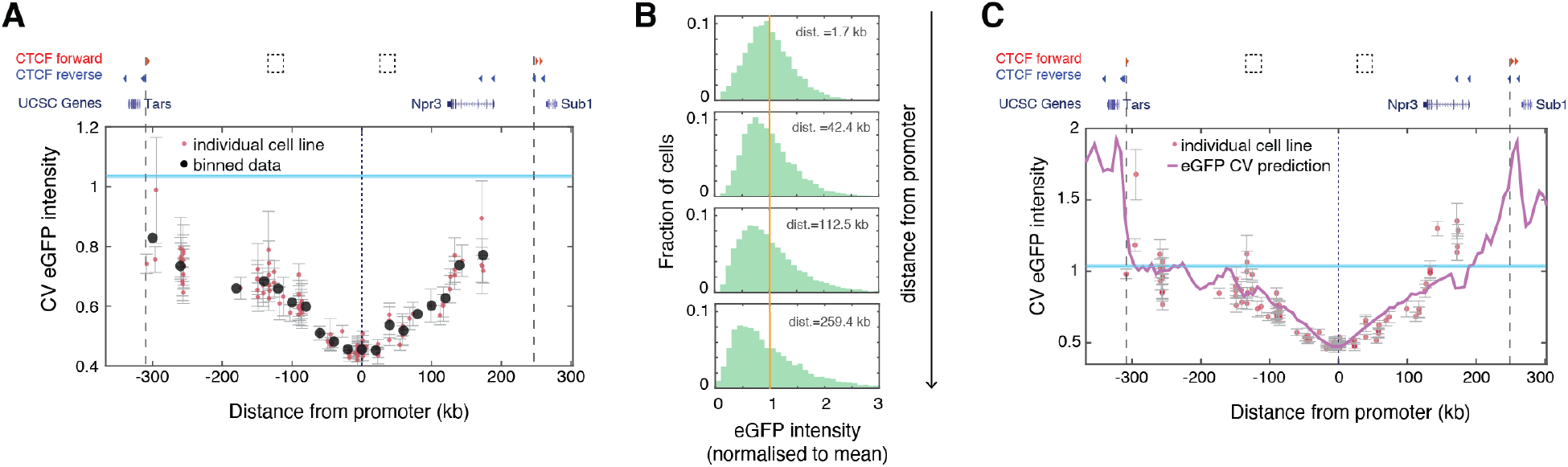
Enhancer genomic distance to the promoter modulates cell-to-cell variability in expression levels. A. Coefficients of variation (CV) of eGFP levels measured by flow cytometry plotted against SCR insertion locations in eGFP+ cell lines (light red dots), and averages in 20-kb genomic bins (black dots). Error bars: standard deviation of CV values over three measurements in three different days. Shaded light blue area indicates the interval between mean +/- standard deviation of eGFP level CVs in three promoter-only cell lines. B. Representative eGFP distributions (normalised to mean eGFP level) in clones with increasing absolute genomic distance (1.7 kb, 42.4 kb, 112.5 kb, and 259.43 kb) between the mobilised enhancer and the ectopic Sox2 promoter. Vertical line indicates normalised mean eGFP levels. C. Model prediction (purple line) for eGFP CVs based on the linear conversion of mRNA CVs (Suppl. Fig. 4C). Shaded light blue area indicates the interval between mean +/- standard deviation of eGFP level CVs in three promoter-only cell lines.

Variability in eGFP signals reflects both intrinsic heterogeneity due to the stochastic molecular processes involved in transcription and translation, and extrinsic, cell-context-dependent features that determine transcript and protein content in single cells. Since cell size and cell-cycle phase heterogeneity are major sources of extrinsic variability in transcript and protein abundance^43,44^, we used an established size-gating approach^45,46^ to study how CVs of eGFP distributions in a clonal population vary by isolating cells with progressively more similar size and granularity based on forward and side scatter signals (**Suppl. Fig. 4A**). We found that CVs were only poorly sensitive to increasingly stringent gating (**Suppl. Fig. 4B**). This suggests that intrinsic variation accounts for a major fraction of the observed variability thus indicating that cell-to-cell differences in eGFP levels mostly reflect intrinsic transcriptional heterogeneity. CVs predicted by the model for mRNA distributions were indeed linearly correlated with experimentally observed eGFP CVs (**Suppl. Fig. 4C**), and highly predictive of their dependence on the genomic distance between the SCR and the ectopic *Sox2* promoter (**Fig. 3C**). We conclude that the genomic distance between an enhancer and a promoter is a major determinant of cell-to-cell transcriptional heterogeneity.

Our data provide evidence that in a chromatin environment devoid of structural and regulatory perturbations, transcription from a promoter is determined by a nonlinear transformation of its contact probabilities with an enhancer. To explore if CTCF binding affects this relationship and quantitatively test the transcriptional consequences of introducing a CTCF loop within a TAD, we performed the enhancer mobilisation assay in mESCs where only one of the two internal CTCF sites was homozygously deleted. The remaining forward CTCF site is located 36 kb downstream of the transgene and loops onto the reverse CTCF sites located towards the 3’ end of the domain (**Fig. 4A**). Mobilisation of the SCR in this context resulted in the generation of 172 additional cell lines whose eGFP transcription levels were indistinguishable from those generated in the ‘empty’ TAD, except across the CTCF site that was able to severely, but not completely, insulate the ectopic *Sox2* promoter from the enhancer (**Fig. 4B**). Interestingly, transcriptional insulation provided by the CTCF site did not depend on the absolute distance between the SCR position and the CTCF site itself, with transcription levels across the CTCF site being approximately 60% lower than those generated by the SCR at comparable distances in the absence of the CTCF site (**Fig. 4C**).

**Figure 4.**
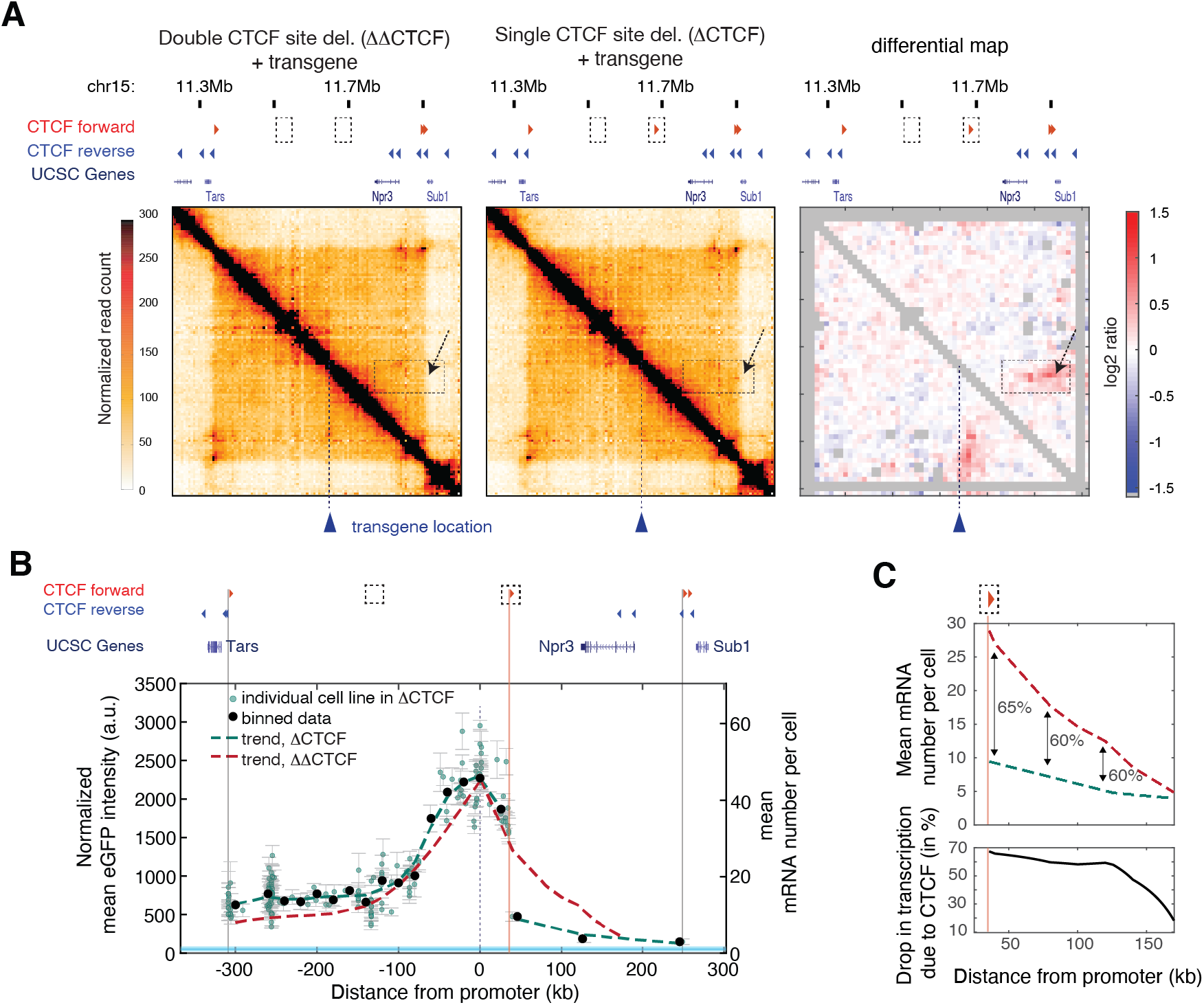
Insulation by a single CTCF site exceeds contact probability changes. A. Capture Hi-C maps (6.4-kb resolution) of founder mESC lines in the absence (left, ΔΔCTCF) or presence (center, ΔCTCF) of a forward CTCF motif 36 kb downstream to the ectopic *Sox2* promoter. Right: differential contact map; grey pixels: correspond to ‘noisy’ interactions that did not satisfy our quality control filters (see Methods). The position of the CTCF site as well as the changes in structure it generates are highlighted by dotted boxes and arrows. B. Mean eGFP levels in individual GFP+ cell lines (green dots, error bars: standard deviation over three measurements in three different days), binned data (black dots), and data trend (green dashed line) after SCR mobilization in the ΔCTCF background (CTCF binding site at +36Kb is highlighted by a vertical pink line). The trend of eGFP levels in individual GFP+ cell lines in the ΔΔCTCF background (red dashed line, Fig. 1H) is shown for comparison. Mean mRNA numbers per cell were inferred from eGFP counts using calibration with smRNA FISH, see Suppl. Fig. 1H. Shaded light blue area indicates the interval between mean +/- standard deviation of eGFP levels in three promoter-only cell lines. C. Upper panel: Zoom-in of the relative decrease in transcription levels measured in GFP+ cell lines in absence (red dashed line) and in presence (dark green dashed line) of the CTCF binding site at +36 kb (vertical pink line). Numbers indicates percent ratios between ΔCTCF and ΔΔCTCF trendlines. Lower panel: percent ratios as a function of distance from the promoter.

Strikingly, transcriptional insulation across the CTCF site occurred in the absence of detectable changes in the promoter’s interaction probabilities with the region downstream of the CTCF site (**Fig. 4A** right). This suggests that a single CTCF site can exert transcriptional insulation through additional mechanisms beyond simply driving physical insulation, possibly depending on site identity^47^ and flanking sequences^18^.

The SCR is a strong enhancer accounting for most of the transcriptional activity of the endogenous *Sox2* gene^31,32^. We reasoned that a weaker enhancer should in principle lead to a different transcriptional response in relation to its contact probabilities with the promoter. We thus challenged the model to predict the outcome of an additional experiment where we modify enhancer strength. There are two ways in which model parameters might change when the strength of the enhancer is reduced. The ratio between transition rates through regulatory steps *k_forward_* and *k_back_* might decrease, resulting in a slower transmission of regulatory information to the promoter (**Fig. 5A**). This would generate a transcriptional response with maximal transcriptional levels that are similar to those generated by the SCR but with different sensitivity to changes in contact probabilities (**Fig. 5A**). Alternatively (although not exclusively) some of the parameters of the high two-state promoter regime could be modified, such as *k_on_* and *k_off_* (**Fig. 5B**). This would conserve the shape of the transcriptional response but decrease maximal transcription levels (**Fig. 5B**).

**Figure 5.**
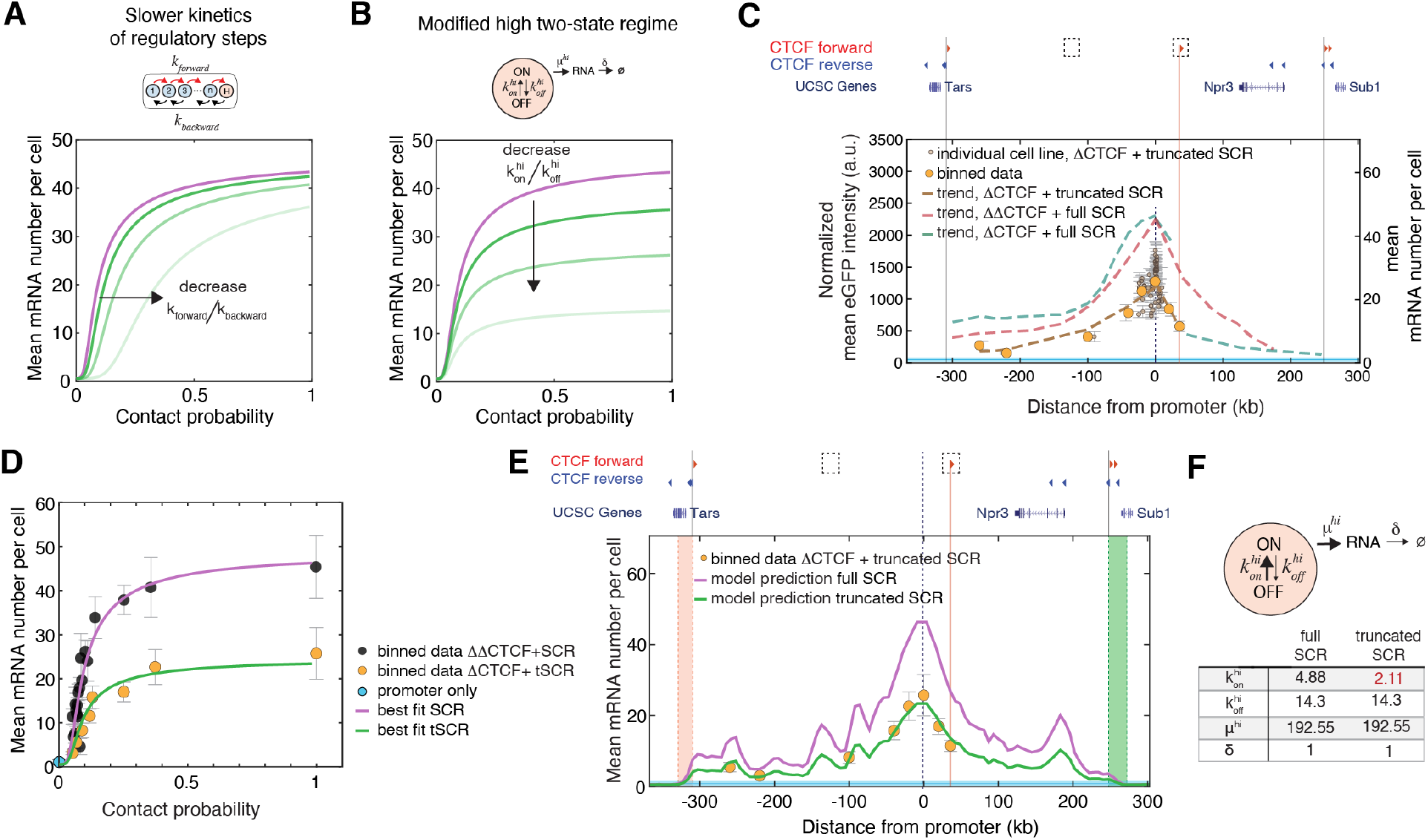
Enhancer strength modulates *on* rates at the promoter and determines sensitivity to insulation through a CTCF site. A. Model predictions under the hypothesis that decreasing enhancer strength results in a slower flow of regulatory information to the promoter, i.e. decreases the ratio between forward and backward rates through intermediate regulatory steps. B. Same as A under the alternative hypothesis that decreasing enhancer strength modifies promoter parameters in the high two-state regime (cf. Fig. 2). C. Mobilisation of the truncated version SCR (tSCR). eGFP levels in individual eGFP+ lines are plotted as a function of the tSCR genomic position. Data from 74 individual cell lines from a single experiment (brown dots; error bars: standard deviation of three measurements performed in three different days) and the average eGFP values calculated within equally spaced 20-kb bins (orange dots) are shown. Dashed brown line: trendline based on a smoothing spline interpolant of the average eGFP values from the indicated experiments. The trends of eGFP levels in individual GFP+ cell lines where the SCR was mobilized either in the ΔΔCTCF background (red dashed line, Fig. 1H) or in the ΔCTCF background (green dashed line, Fig. 4B) are shown for comparison. Mean mRNA numbers per cell were inferred from eGFP counts using calibration with smRNA FISH, see Suppl. Fig. 1H. Shaded light blue area indicates the interval between mean +/- standard deviation of eGFP levels in three promoter-only cell lines. D. The transcriptional response of the truncated SCR (green line) can be predicted from the best fit to the full-length SCR (purple line) with a modified on rate in the high two-state regime (*k^hi^_on_*). E. Model predictions of the eGFP levels generated by the truncated SCR (green line) plotted against binned data from panel C (orange dots). Model prediction for the full-length SCR (purple line) is plotted for comparison. Shaded light blue area indicates the interval between mean +/- standard deviation of eGFP levels in three promoter-only cell lines. Domains across TAD boundaries are highlighted by red and green shaded areas and defined as in Figure 2A. F. Parameters of the high two-state regime for the full-length SCR and truncated SCR (all the other parameters are set to their best-fit values for the full-length SCR, as in Suppl. Fig. 3A).

To test these predictions, we performed the enhancer mobilisation assay but now using a truncated version of the SCR (**Suppl. Fig. 5A**). This version contains only one of the two ~1.5-kb SCR subregions that share similar transcription factor binding sites^31^ and independently operate as weaker enhancers of the *Sox2* promoter in transient reporter assays (**Suppl. Fig. 5B**), as previously reported^31^. Mobilisation of the truncated SCR in mESCs with the forward CTCF site downstream of the promoter (cf. **Fig. 4A**) led to 74 eGFP+ cell lines with approximately 2-fold lower transcription levels than those generated by the full-length enhancer at comparable genomic distances (**Fig. 5C**). Interestingly, contrary to the full-length SCR, the truncated enhancer was completely insulated from the ectopic *Sox2* promoter by the CTCF site (**Fig. 5C**). This shows that the level of functional insulation generated by the same CTCF site critically depends on the strength of the enhancer.

Consistent with model predictions, the transcriptional response generated by the truncated SCR when it was inserted upstream of the CTCF site was also highly nonlinear (**Fig. 5D**). The transcriptional response was in quantitative agreement with model predictions under the hypothesis that enhancer strength modifies the parameters of the high promoter regime rather than those of intermediate regulatory steps (**Fig. 5B**). Both the transcriptional response to contact probabilities (**Fig. 5D**) and the distance dependence of transcription inside the TAD (**Fig. 5E**) could be indeed predicted using the full-length SCR best-fit parameters with a decreased *on* rate in the high promoter regime (**Fig. 5F**). This further strengthens our interpretation of the data in terms of the model and implies that enhancer strength modulates the ability of a promoter to turn *on*, with a substantial impact on burst frequency but not on burst size (**Suppl. Fig. 5CD**).

In the nonlinear transcriptional response we identified, high sensitivity to changes in the low contact probability regime (i.e. at long genomic distances) allows to secure complete insulation by TAD boundaries of even strong enhancers such as the full-length SCR. Interestingly, the majority (~75%) of active promoters in mESCs have contact probabilities with the nearest TAD boundary that are comparable to those in our experiments (lower than 0.2) (**Suppl. Fig. 5E**, shaded area). These promoters should therefore experience the same insulation mechanism. A fraction of promoters are however closer (or directly adjacent) to a TAD boundary and thus perceive the boundary at larger contact probabilities, where the transcriptional response is less sensitive to changes in contact probabilities (**Suppl. Fig. 5E**). Insulation of these promoters might at least in some cases be ensured by the fact that boundaries become stronger (i.e. drops in contact probabilities across the boundary increase) with decreasing distance from the promoter (**Suppl. Fig. 5F**). Boundaries associated with (clusters of) CTCF sites might additionally benefit from the fact that insulation from CTCF sites can exceed the changes in contact probabilities they generate (**Fig. 4**).

In conclusion, our study provides unbiased and systematic measurements of promoter output as a function of a large number of densely located enhancer positions, in the presence of minimal confounding effects. Analysis of hundreds of cell lines allows us to move beyond locus-specific observations and establishes a quantitative framework for understanding the role of chromosome structure in long-range transcriptional regulation.

Our data reveal that within a TAD, absolute transcription levels generated by an enhancer depend on its genomic distance from the promoter and are determined by a nonlinear relationship with their contact probabilities. Minimal regulatory and structural complexities introduce deviations from this behavior and might thus confound its detection outside a highly controlled genomic environment, i.e. when studying regulatory sequences in their endogenous context^25^. Mathematical modeling and validation against experimental data reveals that the observed nonlinear transcriptional response is generated by enhancer interactions being memorized into longer-lived transcriptional states in individual cells. In addition to readily explaining the absence of correlation between transcription and physical proximity in single-cell experiments, this argues that absence of such correlation should not be interpreted as absence of causality in future experiments.

The observed nonlinear transcriptional response provides a potential mechanism for how TAD boundaries can insulate a substantial fraction of promoters. Our experiments however additionally reveal that enhancer strength is not only a determinant of absolute transcription levels, but also of the level of insulation provided by CTCF. Taken together, our data thus imply that transcriptional insulation is not an intrinsic absolute property of all TAD boundaries or CTCF interactions, but rather a graded, context-dependent variable depending on enhancer strength, boundary strength, and distance from a promoter.

## Supporting information

Supplementary Model Description

## METHODS

### Culture of embryonic stem cells

All cell lines are based on E14 mouse embryonic stem cells (mESCs). Cells were cultured on gelatin-coated culture plates in Glasgow Minimum Essential Medium (Sigma-Aldrich, G5154) supplemented with 15% foetal calf serum (Eurobio Abcys), 1% L-Glutamine (Thermo Fisher Scientific, 25030024), 1% Sodium Pyruvate MEM (Thermo Fisher Scientific, 11360039), 1% MEM Non-Essential Amino Acids (Thermo Fisher Scientific, 11140035) 100μM β-mercaptoethanol, 20 U/ml leukemia inhibitory factor (Miltenyi Biotec, premium grade) in 8% CO2 at 37°C. Cells were tested for mycoplasma contamination once a month and no contamination was detected. After piggyBac-enhancer transposition, cells were cultured in standard E14 medium supplemented with 2i (1 uM MEK inhibitor PDO35901 (Axon, 1408) and 3 uM GSK3 inhibitor CHIR 99021 (Axon, 1386)).

### Generation of enhancer-promoter piggyBac targeting vectors

Homology arms necessary for the knock-in, the *Sox2* promoter, the *Sox2* control region (SCR) and the truncated version of the SCR (Ei) were amplified from E14 mESCs genomic DNA by Phusion High-Fidelity DNA Polymerase (Thermo Scientific, F549) using primers compatible with Gibson assembly cloning (NEB, E2611). The targeting vector was generated starting from the 3-SB-EF1-PBBAR-SB plasmid^48^, kindly gifted by Rob Mitra. To clone homology arms into the vector, BspEI and BclI restrictions sites were introduced using Q5^®^ Site-Directed Mutagenesis Kit (NEB, E0554). The left homology arm was cloned using Gibson assembly strategy by linearizing the vector with BspEI (NEB, R0540). The right homology arm was cloned using Gibson assembly strategy by linearizing the vector with BclI (NEB, R0160). The *Sox2* promoter was cloned by first removing the Ef1a promoter from the 3-SB-EF1-PBBAR-SB vector using NdeI (NEB, R0111) and SalI (NEB, R0138) and subsequently using Gibson assembly strategy. The SCR and its truncated version (tSCR or Ei) were cloned between the piggyBac transposon-specific inverted terminal repeat sequences (ITR) by linearizing the vector with BamHI (NEB, R3136) and NheI (NEB, R3131). A transcriptional pause sequence from the human alpha2 globin gene and a SV40 polyA sequence were inserted at both 5’ and 3’ end of the enhancers using Gibson assembly strategy. A selection cassette carrying the Puromycin resistance gene driven by the PGK promoter and flanked by FRT sites was cloned in front of the *Sox2* promoter by linearizing the piggyBac vector with the AsiSI (NEB, R0630) restriction enzyme. Primers used for cloning are listed in **Supplementary_Table_S1**.

### Generation of founder mESC lines carrying the piggyBac transgene

The gRNA sequence for the knock-in of the piggyBac transgene on the chromosome 15 was designed using the online tool https://eu.idtdna.com/site/order/designtool/index/CRISPR_SEQUENCE and purchased from Microsynth AG. gRNA sequence was cloned into the PX459 plasmid (Addgene) using the BsaI restriction site. E14 mESC founder lines carrying the piggyBac transgene were generated using nucleofection with the Amaxa 4D-Nucleofector X-Unit and the P3 Primary Cell 4D-Nucleofector X Kit (Lonza, V4XP-3024 KT). 2×10^6^ cells were harvested with accutase (Sigma Aldrich, A6964) and resuspended in 100 μl transfection solution (82ul primary solution, 18ul supplement, 1μg piggyBac targeting vector carry either the SCR or truncated SCR or promoter alone and 1ug of PX459 ch15_gRNA/Cas9) and transferred in a single Nucleocuvette (Lonza). Nucleofection was performed using the protocol CG110. Transfected cells were directly seeded in pre-warmed 37°C culture in E14 standard medium. 24 hours after transfection, 1ug/mL of puromycin (InvivoGen, ant-pr-1) was added to the medium for 3 days to select cells transfected with PX459 gRNA/Cas9 vector. Cells were then cultured in standard E14 medium for additional 4 days. To select cells with insertion of the piggyBac targeting vector, a second pulse of puromycin was carried out by culturing cells in standard medium supplemented with 1ug/mL of puromycin. After 3 days of selection, single cells were isolated by fluorescence-activated cell sorting (FACS sort) on 96 well-plate. Sorted cells were kept for 2 days in standard E14 medium supplemented by 100 μg/uL primocin (InvivoGen, ant-pm-1) and 10uM ROCK inhibitor (STEMCELL Technologies, Y-27632). Cells were then cultured in standard E14 medium with 1ug/mL of puromycin. Genomic DNA was extracted by lysing cells with lysis buffer (100mM Tris-HCl pH8.0, 5mM EDTA, 0.2% SDS, 50mM NaCl and proteinase K and RNase) and subsequent isopropanol precipitation. Individual cell lines were analyzed by genotyping PCR to determine heterozygous insertion of the piggyBac donor vector. Cell lines showing the corrected genotyping pattern were selected and expanded. Primers used for genotyping are listed in **Supplementary_Table_S1**.

### Puromycin resistance cassette removal

1×10^6^ cells were transfected with 2ug of a pCAG-FlpO-P2A-HygroR plasmid encoding for the flippase (Flp) recombinase using Lipofectamine 3000 (Thermo Fisher Scientific, L3000008) according to the manufacturer’s instructions. Transfected cells were cultured in standard E14 medium for 7 days. Single cells were then isolated by FACS sort on 96 well-plate. Genomic DNA was extracted by lysing cells with lysis buffer (100mM Tris-HCl pH8.0, 5mM EDTA, 0.2% SDS, 50mM NaCl and proteinase K and RNase) and subsequent isopropanol precipitation. Individual cell lines were analyzed by genotyping PCR to verify the deletion of the puromycin resistance cassette. Primers used for genotyping are listed in **Supplementary_Table_S1**. Cell lines showing the correct genotyping pattern were selected and expanded. Selected cell lines were subjected to targeted nanopore sequencing with Cas9-guided adapter ligation (nCATS)^33^ and only the ones showing unique integration of the piggyBac donor vector were used as founder lines for the enhancer mobilization experiments.

### Mobilisation of the piggyBac-enhancer cassette

A mouse codon-optimized version of the piggyBac transposase (PBase) was cloned in frame with the red fluorescent protein tagRFPt (Evrogen) into a pBroad3 vector (pBroad3_hyPBase_IRES_tagRFPt) using Gibson assembly cloning (NEB, E2611). 2×10^5^ cells were transfected with 0.5ug of pBroad3_hyPBase_IRES_tagRFPt using Lipofectamine 3000 (Thermo Fisher Scientific, L3000008) according to the manufacturer’s instructions. To increase the probability of enhancer transposition, typically 12 independent PBase transfections were performed at the same time in 24-well plates. Transfection efficiency as well as expression levels of hyPBase_IRES_tagRFPt transposase within the cells population were monitored by flow cytometry analysis. 7 days after transfection with PBase, individual eGFP+ cell lines were isolated by FACS sort in 96 well-plates. Sorted cells were kept for 2 days in standard E14 medium supplemented by 100 μg/m primocin (InvivoGen, ant-pm-1) and 10uM ROCK inhibitor (STEMCELL Technologies, Y-27632). Cells were cultured in E14 standard medium for additional 7 days and triplicated for genomic DNA extraction, flow cytometry analysis and freezing.

### Sample preparation for mapping piggyBac-enhancer insertion sites in individual cell lines

Mapping of enhancer insertion sites in individual cell lines was performed using splinkerette PCR. The protocol was performed as in Uren et al.^49^ with a small number of modifications. Genomic DNA from individual eGFP+ cell lines was extracted from 96 well-plates using Quick-DNA Universal 96 Kit (Zymo Research, D4071) according to the manufacturer’s instructions. Purified Genomic DNA was digested by 0.5uL of Bsp143I restriction enzyme (Thermo Scientific, FD0784) for 15 minutes at 37°C followed by a heat inactivation step at 65°C for 20 min. Long (HMSpAa) and short (HMSpBb) splinkerette adaptors were first resuspended with 5X NEBuffer 2 (NEB, B7002) to reach a concentration of 50uM. 50uL of HMSpA adapter was then mixed with 50uL of HMSpBb adapter (Aa+Bb) to reach a concentration of 25uM. The adapter mix was denatured and annealed by heating it to 95°C for 5 min and then cooling to room temperature. 25pmol of annealed splinkerette adaptors were ligated to the digested genomic DNA using 5U of T4 DNA ligase (Thermo Fisher, EL0011) and incubating the samples for 1 hour at 22°C followed by a heat inactivation step at 65°C for 10 min. For Splinkerette amplifications, PCR#1 was performed combining 2uL of the splinkerette sample, 1U of Platinum Taq polymerase (Thermo Fisher Scientific, 10966034), 0.1uM of HMSp1 and 0.1uM of PB5-1 (or PB3-1) primer while Splinkerette PCR#2 was performed using 2uL of PCR#, 1U of Platinum Taq polymerase (Thermo Fisher Scientific, 10966034), 0.1uM of HMSp2 and 0.1uM of PB5-5 (or PB3-2) primer. The quality of PCR amplification was checked by Agarose gel. Samples were sent for Sanger Sequencing (Microsynth AG) using the PB5-2 (or PB3-2) primer. Primers used for Splinkerette PCRs and sequencing are listed in **Supplementary_Table_S1**. Mapping of enhancer insertion sites in individual cell lines was performed as described in “*Mapping of piggyBac-enhancer insertion sites in individual cell lines*”.

### Flow cytometry eGFP fluorescence intensities measurements and analysis

eGFP+ cell lines were cultured in Serum+2i medium for two weeks prior to flow cytometry measurements. eGFP levels of individual cell lines were measured on a BD LSRII SORP flow cytometer using BD High Throughput Sampler (HTS), which enabled sample acquisition in 96-well plate format. Measurements were repeated three times for each clone. Mean eGFP fluorescence intensities were calculated for each clone using FlowJo and all three replicates were averaged.

### Normalisation of mean eGFP fluorescence intensities

Mean eGFP fluorescence levels of each cell line measured in flow cytometry were first corrected by subtracting the mean eGFP fluorescence intensities measured in wild-type E14 mESC cultured in the same 96-well plate. The resulting mean intensities were then normalized by dividing them by the average mean intensities of all cell lines where the SCR was located within a 40kb window centered at the promoter location, and multiplied by a common factor.

### Sample preparation for high-throughput sequencing of piggyBac-enhancer insertion sites

5×10^5^ cells were transfected with 2ug of PBase using Lipofectamine 3000 (Thermo Fisher Scientific, L3000008) according to the manufacturer’s instructions. Transfection efficiency as well as expression levels of PBase within the cells population were monitored by flow cytometry analysis. 5 days after transfection with PBase, genomic DNA was purified using DNeasy Blood & Tissue Kit (Qiagen, 69504). To reduce the contribution from cells where excision of piggyBac-enhancer did not occur, we depleted eGFP sequences using an *in vitro* Cas9 digestion strategy. gRNAs sequences for eGFP depletion were designed using the online tool https://eu.idtdna.com/site/order/designtool/index/CRISPR_SEQUENCE (**Supplementary_Table_S1**). Custom designed Alt-R CRISPR-Cas9 crRNAs containing the gRNA sequences targeting eGFP (gRNA_1_3PRIME and gRNA_2_3PRIME), Alt-R CRISPR-Cas9 tracrRNA (IDT, 1072532) and Alt-R S.p. Cas9 enzyme (IDT, 1081060) were purchased from IDT. In vitro cleavage of the eGFP fragment by Cas9 was performed following the IDT protocol “In vitro cleavage of target DNA with ribonucleoprotein complex”. In brief, 100 μM of Alt-R CRISPR-Cas9 crRNA and 100 μM of Alt-R CRISPR-Cas9 tracrRNA were assembled by heating the duplex at 95°C for 5 minutes and allowing to cool to room temperature (15–25°C). To assemble the RNP complex, 10 μM of Alt-R guide RNA (crRNA:tracrRNA) and 10 μM of Alt-R S.p. Cas9 enzyme were incubated at RT for 45 minutes. To perform in vitro digestion of eGFP, 300ng of genomic DNA extracted from the pool cells transfected with the PBase were incubated for 2 hours with 1 μM Cas9/RNP. After the digestion, 40ug of Proteinase K were added and the digested sample was further incubated at 56°C for 10 min to release the DNA substrate from the Cas9 endonuclease. After purification using AMPURE beads XP (Beckman Coulter, A63881), genomic DNA was digested by 0.5uL of Bsp143I restriction enzyme (Thermo Scientific, FD0784) for 15 minutes at 37 °C followed by a heat inactivation step at 65°C for 20 min. 125pmol of annealed splinkerette adaptors (Aa+Bb) were then ligated to the digested genomic DNA using 30U of T4 DNA ligase HC (Thermo Fisher, EL0013) and incubating the samples for 1 hour at 22 °C followed by a heat inactivation step at 65°C for 10 min. For splinkerette amplifications, 96 independent PCR#1 were performed combining 100ng of the splinkerette sample, 1U of Platinum Taq polymerase (Thermo Fisher Scientific, 10966034), 0.1uM of HMSp1 and 0.1uM of PB3-1 primer while Splinkerette PCR#2 was performed using 4uL of PCR#1, 1U of Platinum Taq polymerase (Thermo Fisher Scientific, 10966034), 0.1uM of HMSp2 and 0.1uM of PB3-2 primer. Primers used for Splinkerette PCRs are listed in **Supplementary_Table_S1**. Splinkerette amplicon products were processed using the NEB Ultra II kit according to the manufacturer protocol, using 50ng of input material. Mapping of genome wide insertions was performed as described in “*Mapping of piggyBac-enhancer insertion sites in population-based splinkerette PCR*”.

### Deletion of genomic regions containing CTCF binding sites

gRNAs sequences for depletion of the genomic regions containing the CTCF binding sites were designed uisng the online tool https://eu.idtdna.com/site/order/designtool/index/CRISPR_SEQUENCE and purchased from Microsynth AG (**Supplementary_Table_S1**). gRNAs sequences were cloned into the PX459 plasmid (Addgene) using the BsaI restriction site. To remove the first forward CTCF binding site (chr15:11520474-11520491), 3×10^5^ cells were transfected with 0.5ug of PX459 CTCF_KO_gRNA3/Cas9 and 1ug of PX459 CTCF_KO_gRNA10/Cas9 plasmids using Lipofectamine 2000 (Thermo Fisher Scientific, 11668019) according to the manufacturer’s instructions. To remove the second forward CTCF binding sites (chr15:11683162-11683179), 1×10^6^ cells were transfected with 1ug of PX459 gRNA2_CTCF_KO/Cas9 and 1ug of PX459 gRNA6_CTCF_KO/Cas9 plasmids using Lipofectamine 2000 (Thermo Fisher Scientific, 11668019) according to the manufacturer’s instructions. 24 hours after transfection, 1ug/mL of puromycin was added to the medium for 3 days. Cells were then cultured in standard E14 medium for additional 4 days. To select cell lines with homozygous deletion, single cells were isolated by FACS sort on 96 wellplate. Sorted cells were kept for 2 days in E14 standard medium supplemented by 100 μg/m primocin (InvivoGen, ant-pm-1) and 10uM ROCK inhibitor (STEMCELL Technologies, Y-27632). Cells were then cultured in standard E14 medium. Genomic DNA was extracted by lysing cells with lysis buffer (100mM Tris-HCl pH8.0, 5mM EDTA, 0.2% SDS, 50mM NaCl and proteinase K and RNAse) and subsequent isopropanol precipitation. Individual cell lines were analyzed by genotyping PCR to determine homozygous deletion of the genomic regions containing the CTCF binding sites. Cell lines showing the corrected genotyping pattern were selected and expanded. Primers used for genotyping are listed in **Supplementary_Table_S1**.

### Single-molecule RNA FISH

Cells were harvested with accutase (Sigma Aldrich, A6964) and adsorbed on poly-L-Lysine (Sigma, P8920) pre-coated coverslips. Cells were then fixed with 3% PFA (EMS, 15710) in PBS for 10 minutes at RT, washed with PBS and kept in 70% ethanol at −20°C. After at least 24 hours incubation in 70% ethanol, coverslips were incubated for 10min with freshly prepared wash buffer composed of 10% formamide (Millipore Sigma, S4117) in 2xSSC (Sigma Aldrich, S6639). Coverslips were hybridized overnight (~16h) at 37°C in freshly prepared hybridization buffer composed of 10% formamide, 10% dextran sulfate (Sigma Aldrich, D6001) in 2xSSC and containing 125nM of RNA FISH probe sets against *Sox2* labeled with Quasar 670 (Stellaris) and against eGFP labeled with Quasar 570 (Stellaris). After hybridization, coverslips were washed twice with wash buffer pre-warmed to 37°C for 30min at 37°C while shaking, followed by 5min incubation with 500ng/ml DAPI solution (Sigma Aldrich, D9564) in PBS (Sigma Aldrich, D8537). Coverslips were then washed twice in PBS and mounted on slides with Prolong Gold medium (Invitrogen, P36934) and cured at room temperature for 24 hours. The coverslips were then sealed and imaged within 24 hours.

### RNA FISH image acquisition

Images were acquired on a Zeiss Axion Observer Z1 microscope equipped with 100mW 561nm and 100mW 642nm HR diode solid-state lasers, Andor iXion 885 EMCCD camera, and α Plan-Fluar 100x/1.45 Oil objective. Quasar 570 signal was collected with the DsRed ET filter set (AHF analysentechnik, F46-005), Quasar 670 with Cy5 HC mFISH filter set (AHF analysentechnik, F36-760), and DAPI with Sp. Aqua HC-mFISH filter set (AHF analysentechnik, F36-710). Typical exposure time for RNA FISH probes was set to around 300-500ms with 15-20 EM gain and 100% laser intensity. DAPI signal was typically imaged with exposure time of 20ms with EM gain 3 and 50% laser intensity. Pixel size of the images were 0.080×0.080μm with z-step of 0.25μm for around 55-70 z planes.

### Image processing and quantification of mRNA numbers

Raw images were processed in KNIME and python in order to extract numbers of RNAs per cell. The KNIME workflow described below is based on workflow published and described in Voigt et al.^50^. Z-stacks were first projected to a maximal projection for each fluorescent channel. Individual cells were then segmented using the DAPI channel using Gaussian convolution (sigma=3), followed by filtering using global threshold with Otsu filter, watershed, and connected component analysis for nuclei segmentation. Cytoplasmic areas were then estimated with seeded watershed. Cells with nuclei partially outside the frame of view were automatically excluded. Cells containing obvious artifacts, wrongly segmented, or not fully captured in xyz dimensions were manually excluded from the final analysis. Spot detection is based on the laplacian of gaussian (LoG) method implemented in TrackMate^51^. For the channels containing RNA-FISH probes signal, RNAs spots were detected after background subtraction (rolling ball radius 20-25px) by selecting spot size 0.2 um and threshold for spot detection based on visual inspection of multiple representative images. Spot detection is based on the Laplacian of Gaussian method from TrackMate. Subpixel localization of RNA spots were detected for RNA channels and a list of spots per cell for each experimental condition and replicate was generated. Spots in each channel were then aggregated by cell in python to extract the number of RNAs per cell.

### Enhancer reporter assays

To generate vectors for enhancer reporter assay, the *Sox2* promoter, *Sox2* control region (SCR) and the truncated versions of the SCR (Ei and Eii) were amplified from E14 mESCs genomic DNA by Phusion High-Fidelity DNA Polymerase (Thermo Scientific, F549) using primer compatible with gibson assembly strategy. The *Sox2* promoter was cloned in the 3-SB-EF1-PBBAR-SB vector as described before. The SCR and the truncated versions Ei and Eii were cloned in front of the *Sox2* promoter by linearizing the vector with AgeI (NEB, R3552) and subsequently using Gibson assembly cloning. A transcriptional pause sequence from the human alpha2 globin gene and a SV40 polyA sequence were inserted at both 5’ and 3’ end of the enhancers. To test enhancers activity, 3×10^5^ cells were co-transfected with 0.5ug of the different versions piggyBac vectors and 0.5ug of pBroad3_hyPBase_IRES_tagRFPt using Lipofectamine 2000 (Thermo Fisher Scientific, 11668019) according to the manufacturer’s instructions. As a control, only 0.5ug of thepiggyBac vector carrying the Sox2 promoter was transfected. 24 hours after transfection, cells were harvested and analyzed by flow cytometry.

### Capture Hi-C sample preparation

20×10^6^ cells were cross-linked with 1% formaldehyde (EMS, 15710) for 10 minutes at RT and quenched with glycine (final concentration 0.125 M). Cells were lysed in 1M Tris-HCL pH8.0, 5M NaCl and 10% NP40 and complete protease inhibitor (Sigma-Aldrich, 11836170001) and subjected to enzymatic digestion using 1000 units of MboI (NEB, R0147). Digested chromatin was then ligated at 16 °C with 10000 U of T4 DNA ligase (NEB, M0202) in ligase buffer supplemented with 10% Triton-100 (Sigma-Aldrich, T8787) and 240ug of BSA (NEB, B9000). Ligated samples were de-crosslinked with 400 μg Proteinase K (Macherey Nagel, 740506) at 65°C and phenol/chloroform purified. 3C library preparation and target enrichment using a custom-designed collection of 6979 biotinylated RNA “baits” targeting single MboI restriction fragments chr15:10283500-13195800 (mm9) (Supplementary Table S2; Agilent Technologies; designed as in Schoenfleder et al.^52^) were performed following the SureSelectXT Target Enrichment System for Illumina Paired-End Multiplexed Sequencing Library protocol. The only exceptions were the use of 9ug of 3C input material (instead of 3ug) and shear of DNA using Covaris sonication with the following settings: Duty Factor: 10%; Peak Incident Power (PIP): 175; Cycles per Burst: 200; Treatment Time: 480 seconds; Bath Temperature: 4°to 8°C).

### Targeted nanopore sequencing with Cas9-guided adapter ligation (nCATS)

gRNAs sequences targeting specific genomic regions of chromosome 15 external to the homology arms of the transgene were designed using the online tool https://eu.idtdna.com/site/order/designtool/index/CRISPR_SEQUENCE (**Supplementary_Table_S1**). Custom designed Alt-R CRISPR-Cas9 crRNAs (5 crRNAs targeting the region upstream and 5 crRNAs targeting the region downstream the integrated transgene), Alt-R CRISPR-Cas9 tracrRNA (IDT, 1072532) and Alt-R S.p. Cas9 enzyme (IDT, 1081060) were purchased from IDT. Samples preparation and Cas9 enrichment were performed following the protocol previously described^33^ with few modifications. Genomic DNA from mESC founder lines was extracted with Gentra Puregene Cell Kit (Qiagen, 158745) following the manufacturer’s instructions. Quality of the High Molecular Weight (HMW) DNA was checked with the TapeStation (Agilent). Typically 5ug of HMW DNA was subjected to incubation using Shrimp Alkaline Phosphatase (rSAP; NEB, M0371) for 30min at 37°C followed by 5min at 65°C to dephosphorylate DNA free ends. For Cas9 enrichment of the target region, all ten Alt-R CRISPR-Cas9 crRNAs were first pooled in equimolar amount (100uM) and subsequently incubated with 100uM of Alt-R CRISPR-Cas9 tracrRNA at 95°C for 5 minutes to assemble the Alt-R guide RNA duplex (crRNA:tracrRNA). To assemble the RNP complex, 4 pmol of Alt-R S.p Cas9 enzyme were incubated with 8pmol Alt-R guide RNA (crRNA:tracrRNA) at RT for 20 minutes. In vitro digestion and A-tailing of the DNA were performed by adding 10uL of the RNP complex, 10mM of dATP (NEB, N0440) and 5U of Taq Polymerase (NEB, M0267) and incubating the samples at 30min 37°C followed by 5min 72°C. Adaptor ligation for Nanopore sequencing was performed using Ligation Sequencing Kit (Nanopore, SQK-CAS109) according to the manufacturer’s instructions. After purification with AMPure PB beads (Witec AG, 100-265-900), samples were loaded into MniION selecting SQK-CAS109 protocol.

### Nanopore sequencing analysis

To map nanopore sequencing reads, we first built a custom ‘genome’ consisting of the transgene sequence flanked by ~10kb mouse genomic sequence upstream and downstream of the target integration site. The custom genome can be found at https://github.com/zhanyinx/Zuin_Roth_2021/blob/main/Nanopore/cassette/cassette.fa. Reads were mapped to the custom genome using minimap2 (v. 2.17-r941) with “-x map-ont” parameter. Nanopore sequencing analysis has been implemented using Snakemake workflow (v. 3.13.3). Reads were visualized using IGV (v. 2.9.4). Full workflow can be found at https://github.com/zhanyinx/Zuin_Roth_2021.

### Capture Hi-C analysis

Capture Hi-C data were analysed using HiC-Pro^53^ (v. 2.11.4) parameters can be found at https://github.com/zhanyinx/Zuin_Roth_2021). Briefly, read pairs were mapped to the mouse genome (build mm9). Chimeric reads were recovered after recognition of the ligation site. Only unique valid pairs mapping to the target regions were used to build contact maps. Iterative correction (ICE)^54^ was then applied on binned data. The target regions can be found at https://github.com/zhanyinx/Zuin_Roth_2021.

### Differential cHi-C maps

To evaluate the structural perturbation induced by the insertion of the transgene and the mobilisation of the enhancer (ectopic sequences), we accounted for differences in genomic distances due to the presence of the ectopic sequence. In the founder cell line (e.g. SCR_ΔΔCTCF), insertion of the transgene modifies the genomic distance between loci upstream and downstream the insertion site. To account for these differences, we generated distance-normalised cHi-C maps where each entry corresponds to the interaction normalised by the corrected genomic distance between the interacting bins. Outliers (defined using the interquartile rule) or bins with no reported interactions from cHi-C were treated as noise and filtered out. Singletons, defined as the top 0.1 percentile of Z-score, were also filtered out. The Z-score is defined as (obs – exp)/stdev, where obs is the cHi-C signal for a given interaction and exp and stdev are the genome-wide average and standard deviation, respectively, of cHi-C signals at the genomic distance separating the two loci. We then calculated the ratios between distance normalized and noise-filtered cHi-C maps. A bilinear smoothing with a window of 2 bins has been applied to the ratio maps to evaluate the structural perturbation induced by the insertion of the ectopic sequence.

### Chromatin states calling with ChromHMM

Chromatin states were called using ChromHMM^30^ with four states. The list of histone modification datasets used can be found in Supplementary Table S3. States with enrichment in H3K9me3 and H3K27me3 were merged, thus resulting in three chromatin states: active (enriched in H3K27ac, H3K36me3, H3K4me1 and H3K9ac), repressive (enriched in H3K9me3 and H3K27me3) and neutral (no enrichment).

### Mapping of piggyBac-enhancer insertion sites in population-based splinkerette PCR

In order to identify true positive enhancer re-insertion sites, we first filtered out reads containing eGFP fragments. We then kept only read-pairs where one side maps to the ITR sequence and the other side maps to the splinkerette adapter sequence. We mapped separately the ITR/splinkerette sides of the read-pair to the mouse genome (build mm9) using BWA mem^55^ with default parameters. Only integration sites which have more than 20 reads from both ITR and splinkerette sides were kept.

### Mapping of piggyBac-enhancer insertion sites in individual cell lines

To map the enhancer position in individual cell lines, Sanger sequencing (Microsynth) without the adaptor sequences are filtered out. The first 24bp of each read after the adapter are then mapped to the mouse genome (mm9) using vmatchPattern (Biostrings v 2.58.0). The script used to map sanger sequencing can be found at https://github.com/zhanyinx/Zuin_Roth_2021.

### Calibration of the mean number of mRNA per cell with smRNA FISH

A linear model was used to predict the average number of eGFP mRNAs based on the mean eGFP intensity. The model was fitted on 7 data points corresponding to the average number of eGFP mRNAs obtained using single-molecule RNA fluorescent in situ and the mean eGFP intensity obtained by Flow cytometry (see **Suppl. Fig. S1H**, *R*^2^ = 0.9749, *p* < 0.0001).

### Mathematical model and parameter fitting

The enhancer-promoter communication model was fitted simultaneously to the mean eGFP levels measured in individual cell lines, to the distribution of RNA numbers measured by smRNA FISH in a control line where the SCR is absent, and in a cell line where the full-length Sox2 control region (SCR) is adjacent to the promoter. The mean number of mRNA was calculated analytically (see Supplementary Information) and the steady-state distribution of the number of mRNA per cell was approximated numerically. All the rate parameters (*k_far_, k_forward_, k_back_, k^lo^_on_, k^lo^_off_, k^hi^_on_, k^hi^_off_, μ^lo^, μ^hi^*) and the number of regulatory steps (*n*) were fitted. First, only the rate parameters corresponding to the low regime (i.e. *k^lo^_on_, k^lo^_off_, μ*) were fitted to the mRNA FISH distribution of the control line where the SCR is absent. In a second step, all the other rate parameters and the number of regulatory steps were fitted simultaneously to the mean eGFP levels measured in individual cell lines and to the distribution of RNA numbers measured by smRNA FISH in the cell line where the full-length Sox2 control region (SCR) is adjacent to the promoter. The model was also fitted to the binned mean number of mRNA molecules inferred from the eGFP+ cell lines with the truncated version of the SCR (**Fig. 4**). In this case several combinations of free parameters were fitted to the data, keeping the other ones fixed to the best fit values obtained for the full-length SCR data set. The different combinations were ranked by the *R*^2^ of the corresponding model prediction. The mathematical model of enhancer-promoter communication and the fitting procedures are explained in detail in the Supplementary Information.

## Data availability

All cHi-C, Oxford Nanopore and population-based splinkerette PCR sequencing fastq files generated in this study have been uploaded to the Gene Expression Omnibus (GEO) under accession GSE172257. The following public databases were used: BSgenome.Mmusculus.UCSC.mm9.

## Code availability

Custom codes generated in this study are available at: https://github.com/zhanyinx/Zuin_Roth_2021 (cHiC, Nanopore, Insertion mapping); https://github.com/gregroth/Zuin_Roth_2021 (mathematical model)

## ACKNOWLEDGMENTS

We would like to thank Rob Mitra for sharing the piggyBac-splitGFP vector, Alistair Boettiger and Jordan Y. Xiao for stimulating discussions on modeling, Marco Michalski and Simon Andrews for capture Hi-C primer design, Laurent Gelman and Jan Eglinger for help with microscopy and image analysis, Gioacchino Natoli, Dirk Schubeler, Helge Grosshans and Elphege Nora for discussions and comments on the manuscript. J.Z was supported by a Marie Skoldowska-Curie grant (748091 - 3DQuant). Research in the Giorgetti lab is funded by the Novartis Foundation, the European Research Council (ERC) under the European Union’s Horizon 2020 research and innovation (759366, ‘BioMeTre’), and the Swiss National Science Foundation (310030_192642). Research in the Meister lab is supported by the Swiss National Science Foundation (IZCOZ0_189884/31003A_176226).

## AUTHOR CONTRIBUTIONS

LG and JZ conceived and designed the study. JZ, JC, EP, MK, GT performed the experiments. GR wrote and analyzed the mathematical model. HK provided assistance with flow cytometry. JC, PM and SS contributed to setting up nCATS. SS also provided assistance with high-throughput sequence experiments. JZ, JC, JR and PM generated cell lines. GR and YZ analyzed the data, except for flow cytometry and single clone insertion mapping (JZ) and smRNA FISH (EP). LG wrote the paper with GR, JZ and YZ and input from all the authors.

---

**Supplementary Figure 1.**
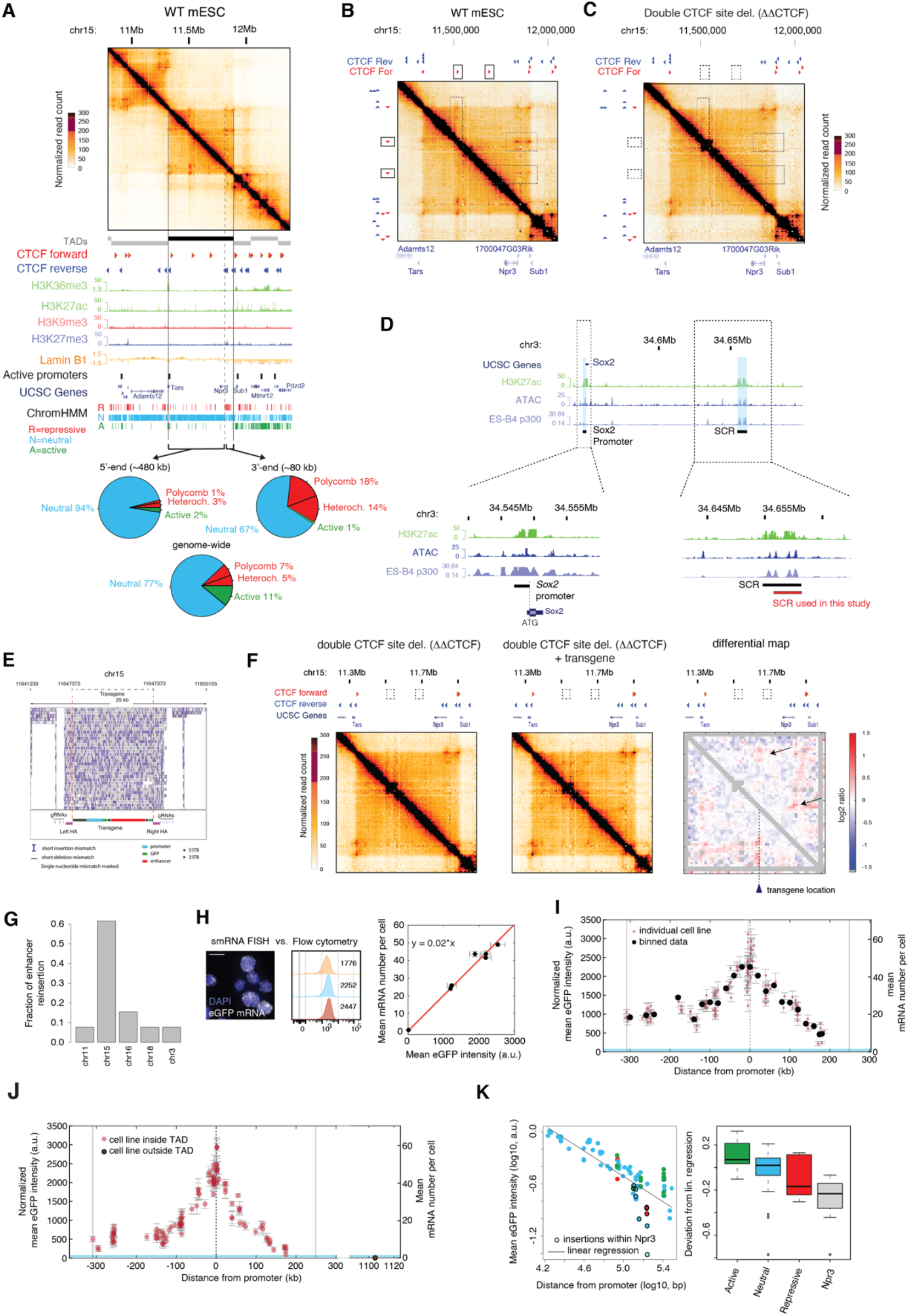
Enhancer action is modulated by genomic distance from the target promoter and constrained by TAD boundaries. A. Top: Capture Hi-C contact map at 6.4-kb resolution in wild-type (WT) mESCs in a 2.6-Mb region centered around the neutral TAD on chromosome 15 we used for the experiments. Vertical grey lines: TAD boundaries. Bottom: genomic datasets and ChromHMM analysis showing that the chosen TAD is devoid of active and repressive chromatin states, with the exception of 80 kb at the 3’-end, which is enriched in repressive chromatin states. B. Close-up view of panel A, highlighting the presence of CTCF-mediated chromatin loops (dotted boxes) in WT mESCs. C. Capture Hi-C contact map at 6.4-kb resolution for the same region as panel B in the cell line with double CTCF site deletions. CTCF deletions lead to loss of CTCF-mediated chromatin loops (dotted boxes). D. Top: UCSC snapshot of the endogenous *Sox2* locus and *Sox2* control region (SCR). Bottom: close-up views showing the regions of the *Sox2* promoter, the SCR region found in ref. 34 and the SCR used in the transgene construct. E. IGV snapshot showing nanopore sequencing reads mapped to a modified mouse genome including the transgene integration. Reads spanning from genomic DNA upstream the left homology arm to genomic DNA downstream the right homology arm confirmed single insertion of the transgene. F. Capture Hi-C maps at 6.4-kb resolution of the mESC line with double CTCF sites deletion (left) and the founder mESC line with transgene insertion (center). Right: differential contact map. Grey pixels correspond to ‘noisy’ interactions that did not satisfy our quality control filters (see Methods). Transgene insertion induces new mild interactions with CTCF sites at the 3’ and 5’ extremities of the TAD (arrows). G. Barplot showing the fraction of piggyBac-SCR reinsertions genome-wide determined by Illumina sequencing of Splinkerette PCR products from a pool of cells after PBase expression. See Methods for a detailed description of the protocol. H. Left: Representative smRNA FISH image and flow cytometry profiles over different passages in a cell line where the SCR was mobilized in the immediate vicinity of the ectopic *Sox2* promoter. Right: Linear relationship between the mean eGFP intensity and the average number of eGFP mRNAs measured using smRNA FISH for seven single cell lines (*R*^2^ = 0.9749, *p* < 0.0001). I. Normalized mean eGFP intensities levels in individual eGFP+ cell lines are plotted as a function of the genomic position of the SCR in individual eGFP+ lines. Data from 128 individual cell lines from a single experiment (light red dots; error bars: standard deviation of three measurements performed in three different days, as in Fig. 1G) and the average eGFP values calculated within equally spaced 20-kb bins (black dots) are shown. Mean mRNA numbers per cell were inferred from eGFP counts using calibration with smRNA FISH, see Suppl. Fig. 1H. Shaded light blue area indicates the interval between mean +/- standard deviation of eGFP levels in three promoter-only cell lines. J. Same plot as Figure 1H including the only SCR insertion we detected outside the TAD boundaries (brown dot). K. Left: Log10 average eGFP expression (from Fig. 1H) as a function of log10 absolute genomic distance between transgene position and SCR reinsertion. Points are color-coded as in panel A (chromHMM active, neutral, and repressive states). Black line denotes linear regression. Black circles denote SCR reinsertions within the Npr3 gene body. Right: deviations of eGFP expression levels from the linear regression correlate with chromatin states called using ChromHMM. Reinsertion of SCR within active or repressive regions respectively increases or decreases enhancer activity compared to neutral regions. Box plot: centre line denotes the median; boxes denote lower and upper quartiles (Q1 and Q3, respectively); whiskers denote 1.5x the interquartile region (IQR) below Q1 and above Q3; points denote outliers.

**Supplementary Figure 2.**
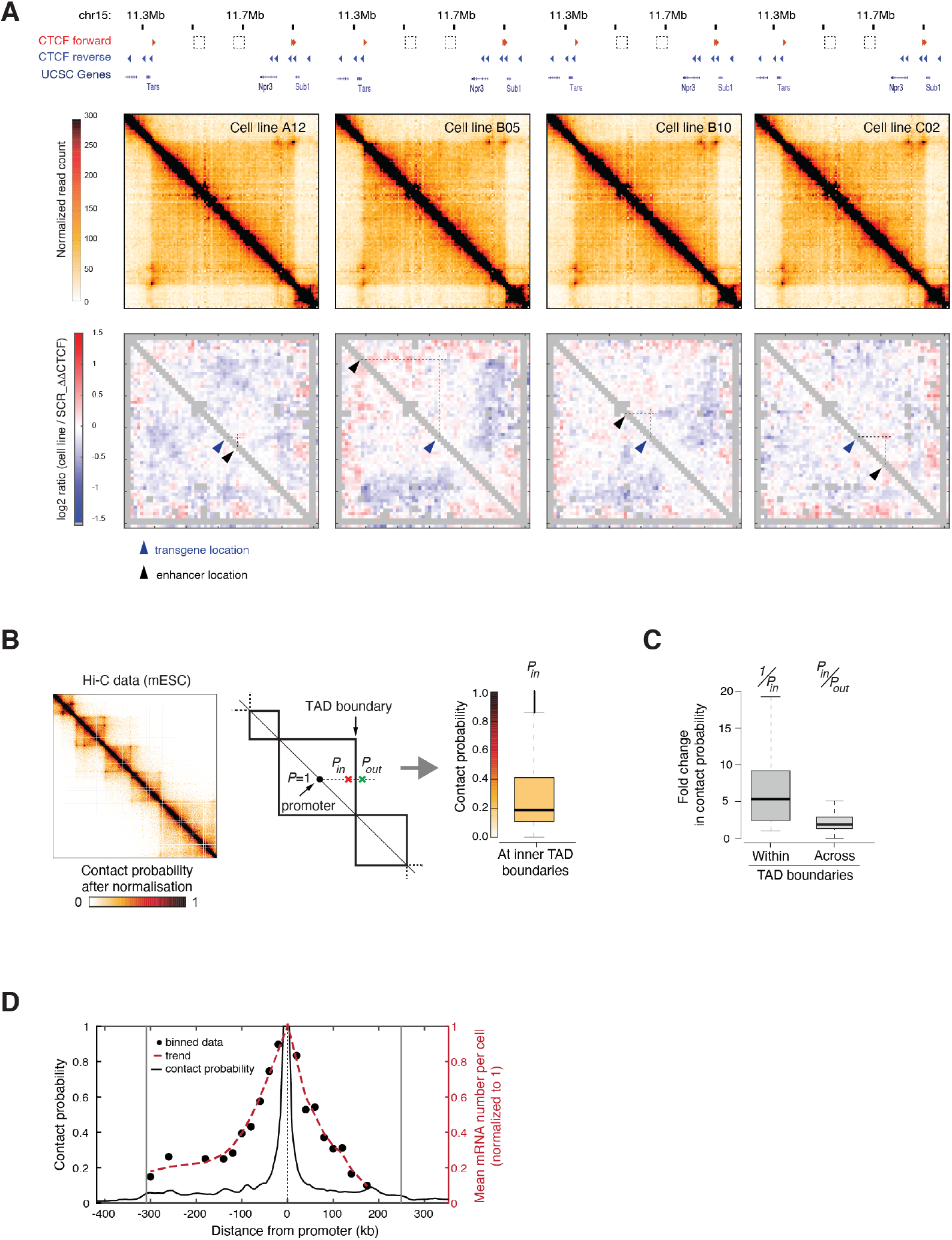
Analysis of chromosome structure around the transgenic locus and genome-wide in mESC. A. Top: Capture Hi-C maps at 6.4-kb resolution of four cell lines where the SCR has been reinserted at different distances (black arrow) from the promoter (blue arrow). Bottom: differential contact map between individual cell lines and the founder line. Grey pixels: correspond to ‘noisy’ interactions that did not satisfy quality control filters (see Methods). B. Left: example of Hi-C heatmap in mESCs at 6.4 kb. Center: scheme depicting how the probability of interaction between a promoter and the region immediately before the nearest TAD boundary (*P_in_*, 12.8 kb i.e. two 6.4-kb bins before the boundary called using CaTCH56) and after the nearest TAD boundary (*P_out_*) are calculated. Right: distribution of contact probability between all active promoters in mESCs and the closest inner TAD boundary (*P_in_*). Box plot description as in Fig. S1K. C. Box plots showing the distribution of contact probability changes within the TAD and across the closest TADs boundary for all active promoters in mESC. Box plot description as in Fig. S1K, outliers not shown. D. Contact probabilities of the founder line from the location of the ectopic *Sox2* transgene (black line) and normalized averaged mean number of mRNAs per cell (highest value =1) generated in individual eGFP+ lines by the SCR mobilization are plotted as a function of its genomic position (dashed red line). The average is calculated within equally spaced 20-kb bins as in Fig. 1H (black dots).

**Supplementary Figure 3.**
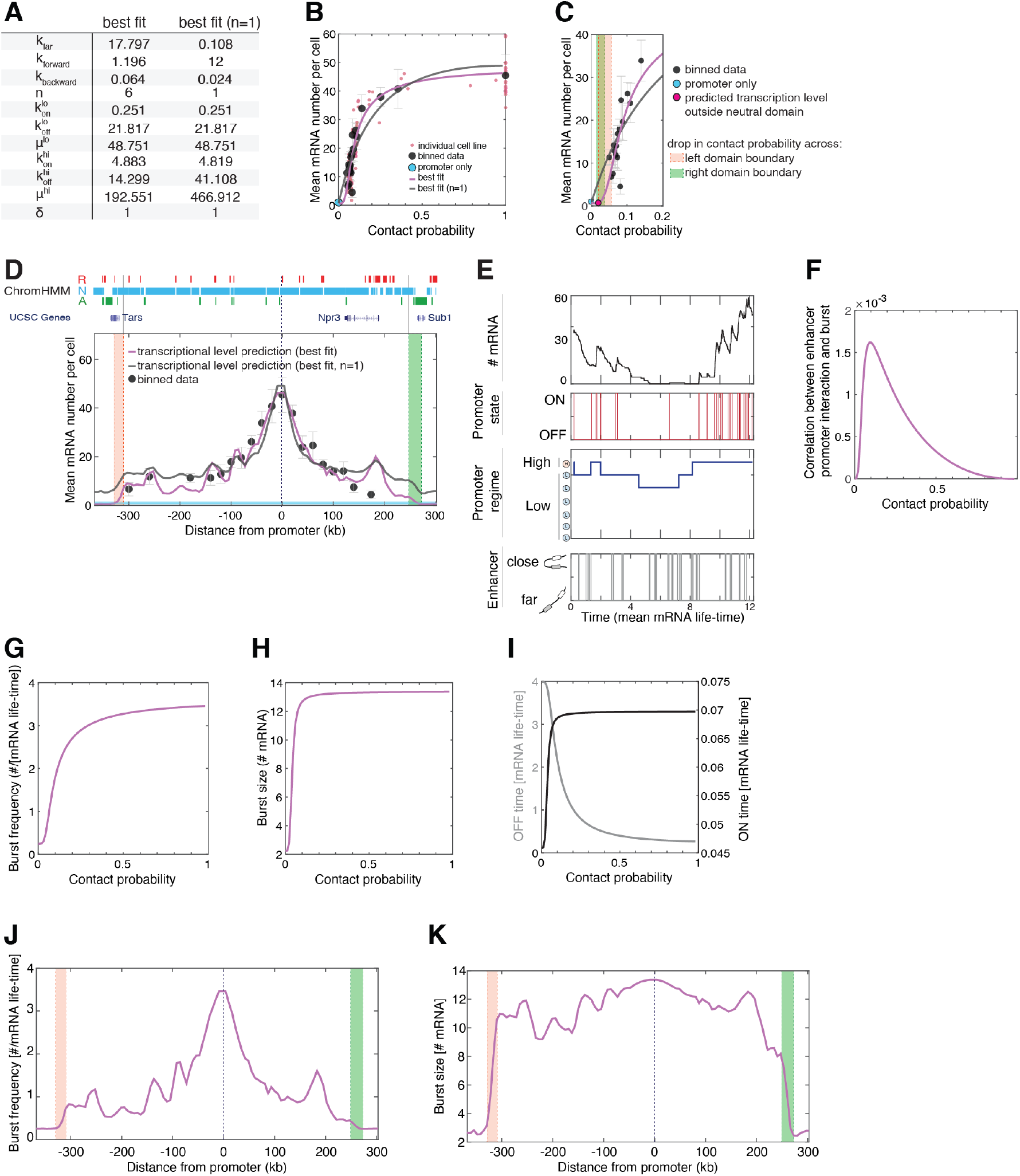
Model fitting and characterization. A. Best fit parameters for the model without constraint on the number of intermediate steps (left column) and for the model constrained to have a single regulatory step (right column). All the rates are expressed in units of the eGFP mRNA life-time. B. Best fit to the experimental data of Fig. 1 panel H when the model is constrained to have a single intermediate step (dark grey line, *R*^2^ = 0.8368) compared to when it is not constrained (purple line, six steps. *R*^2^ = 0.8968). C. Close-up view of panel B highlighting model behavior at low contact probabilities. The model with one intermediate step fails to predict promoter insulation from the SCR outside TAD boundaries. Domains across TAD boundaries are highlighted by red and green shaded areas and defined as in Figure 2A. D. Prediction of transcription levels generated by the SCR within the TAD for both the model with one intermediate step (dark grey line) and the model with six intermediate steps (purple line). The predictions are plotted against binned expression data (black dots; cf. Fig. 1H). Domains across TAD boundaries are highlighted by red and green shaded areas and defined as in Figure 2A. E. Representative single-cell dynamics of enhancer, promoter and RNA states predicted by the model with six intermediate regulatory steps (time unit: mean mRNA life-time). F. Prediction of the correlation between enhancer-promoter interactions (close state) and promoter bursting (ON state) as a function of steady-state contact probability (best fit parameters shown in the left column of panel A). G. Prediction of burst frequency (mean number of bursts per mRNA life-time) as a function of contact probability (best fit parameters shown in the left column of panel A). H. Prediction of burst size (mean number of mRNAs produced per burst) as a function of contact probability (best fit parameters shown in the left column of panel A). I. Prediction of the mean time that the promoter spends in the OFF state (gray line) or the ON state (black line) as a function of contact probability (best fit parameters shown in the left column of panel A). J. Prediction of the burst frequency generated by the SCR within the TAD (best fit parameters shown in the left column of panel A). Domains across TAD boundaries are highlighted by red and green shaded areas and defined as in Figure 2A. K. Prediction of the burst size generated by the SCR within the TAD (best fit parameters shown in the left column of panel A). Domains across TAD boundaries are highlighted by red and green shaded areas and defined as in Figure 2A.

**Supplementary Figure 4.**
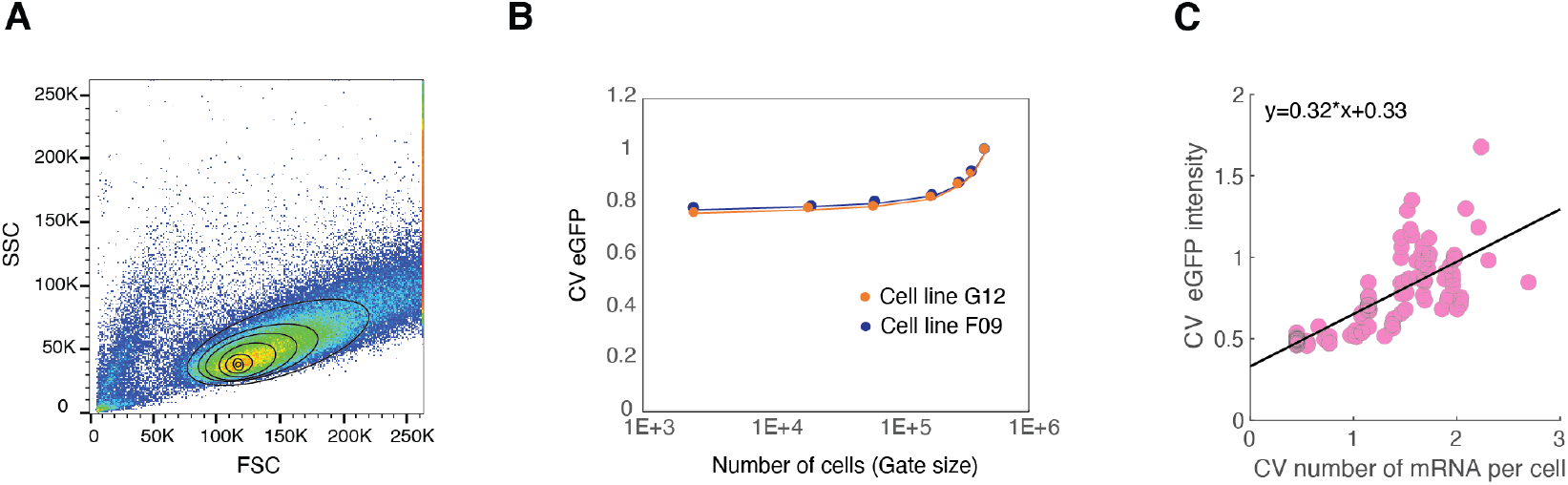
Size-gating approach to estimate the contribution of intrinsic and extrinsic sources of cell-to-cell variability. A. Flow cytometry Forward (FSC) and Side Scatter (SSC) plot representing the gating strategy used to calculate the coefficient of variation of eGFP distribution. Cells are isolated using progressively stringent gates (black circular line). Flow cytometry data are shown for one eGFP+ cell line where the SCR was remobilised within the TAD. B. Coefficient of variation (standard deviation/mean eGFP level) is plotted as a function of cell numbers within each gate size for two eGFP+ cell lines. C. Linear relationship between the coefficients of variation of the eGFP intensities (calculated in individual cell lines as in Figure 3A, light red dots) and coefficients of variation of the number of eGFP mRNAs predicted by the model with the best fit parameters (shown in Fig. 2H) (*R*^2^ = 0.594, *p* < 0.0001).

**Supplementary Figure 5.**
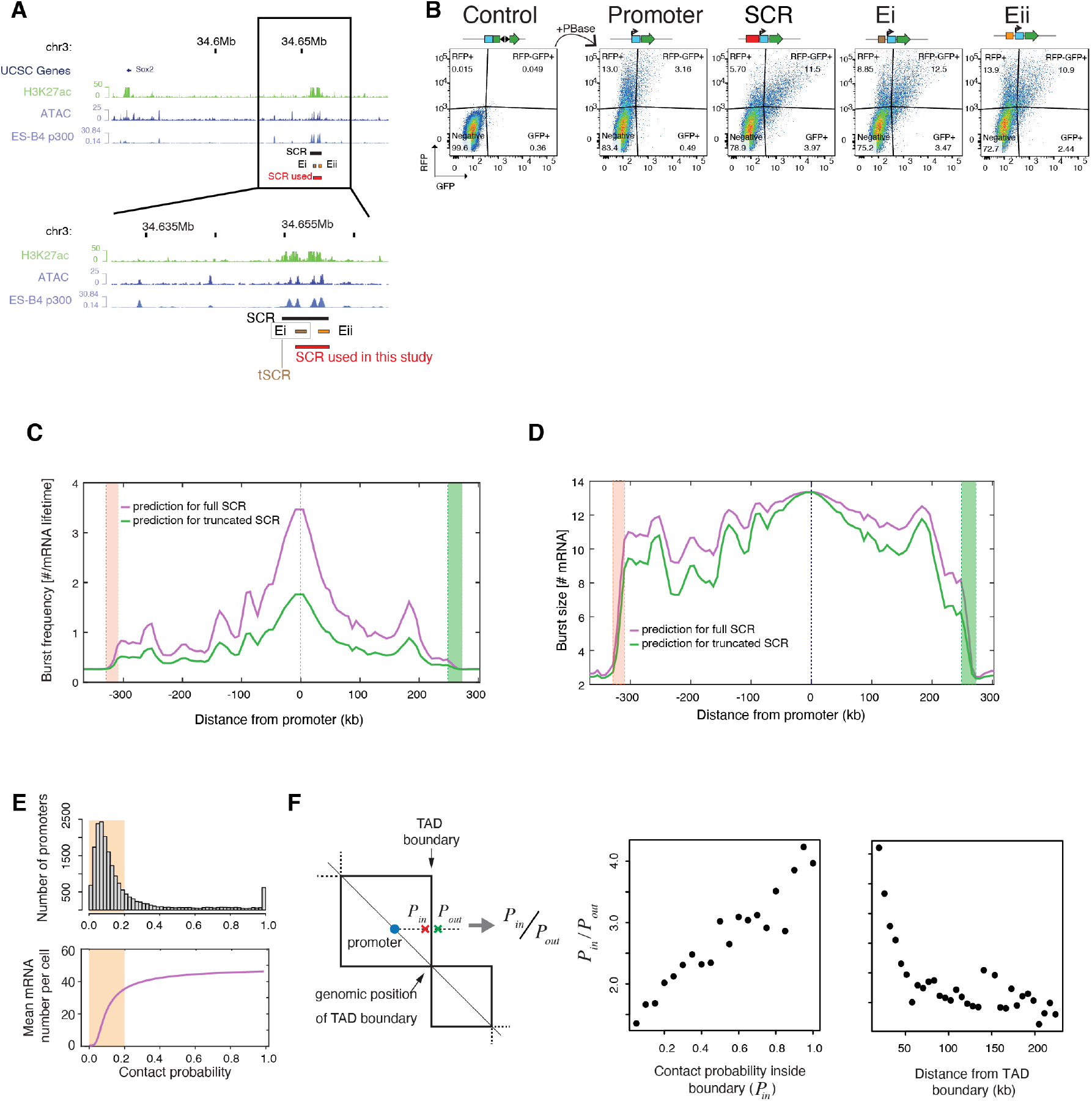
Analysis of truncated SCR and analysis of TAD boundaries genome-wide. A. Top: UCSC genome browser snapshot of the endogenous *Sox2* locus and *Sox2* control region (SCR). Bottom: close-up view showing the SCR (black) identified in ref31 and the enhancer regions used in the transient reporter assays shown in panel B. Full-length enhancer is in red (same as in Figure 1); truncated versions are in brown (Ei) and orange (Eii). Experiments in Figure 5 were performed with Ei. B. Flow cytometry analysis of mESCs transiently transfected with different versions of split eGFP plasmids and PBase-RFP. Split eGFP constructs carry either no enhancer, or the full-length SCR (red, see panel A), or the first (brown-Ei) or second (orange-Eii) SCR subregions in front of the *Sox2* promoter. Transcription levels generated upon co-transfection with PBase are higher in the presence of the full-length SCR compared to truncated versions. Numbers in each quadrant represent the % of cells either negative or RFP, GFP and RFP-GFP positive. C. Prediction of burst frequency generated within the TAD by the full-length SCR (purple line) and the truncated SCR (green line). Parameters: best fit parameters for the full-length SCR as shown in the left column of Fig. S3A, and best fit parameters for the truncated SCR as shown in Fig. 5F. Domains across TAD boundaries are highlighted by red and green shaded areas and defined as in Figure 2A. D. Prediction of the burst size generated within the TAD by the full-length SCR (purple line) and the truncated SCR (green line). Parameters: best fit parameters for the full-length SCR as shown in the left column of Fig. S3A, and best fit parameters for the truncated SCR as shown in Fig. 5F. Domains across TAD boundaries are highlighted by red and green shaded areas and defined as in Figure 2A. E. Top panel: distribution of contact probabilities between all active promoters in mESCs and the nearest inner TAD boundaries, calculated as in Suppl. Fig. 2B. Bottom panel: Model prediction for the mean eGFP mRNA numbers per cell plotted against contact probabilities shown as comparison (same as Fig. 2I). Shaded areas correspond to promoters with contact probability with the closest TAD boundary below 0.2. F. Left panel: scheme of how the probabilities of interaction between promoter and the region before (*P_in_*) and after the TAD boundary (*P_out_*) are calculated, same criteria as in Suppl. Fig. 2B. Central panel: promoters with higher contact probabilities with TAD boundaries experience stronger drops of contact probability across boundaries. Right panel: promoters closer to TAD boundaries experience a stronger drop of contact probability across boundaries.

