## Supplementary Model Description for "Nonlinear control of transcription through enhancer-promoter interactions"

### Supplementary Model description: Nonlinear control of transcription through enhancer-promoter interactions

This Supplementary Information is organised as follows. In Section 1, we describe the mathematical model of enhancer-promoter communication and provide a detailed scheme. In Sections 2 and 3, we calculate from the model the mean and variance of the number of RNAs as a function of the contact probability between the enhancer and the promoter. In Section 4, we calculate from the model the transcriptional burst size and burst frequency. In Section 5, we calculate the correlation between enhancer-promoter interactions and promoter transcriptional activity. In Section 6, we calculate an approximation of the model steady-state distribution of the number of RNAs. In Section 7, we study qualitatively the curve describing the transformation of contact probabilities into mean number of RNAs, and how its shape depends on model parameters. Finally in Section 8, we describe model fitting to the experimental data.

#### 1 Description of the mathematical model of enhancer-promoter communication

The model is fully stochastic and describes the time evolution of four variables: the enhancer state, the promoter state, the promoter regime state, and the number of RNA molecules per cell. The enhancer states represent the relative position of the enhancer and the promoter ($e_1$ := close: the enhancer is in physical proximity of the promoter, i.e. their distance is smaller than an arbitrary threshold; $e_2$ := far: the enhancer is not in physical proximity of the promoter). The promoter states describe the transcriptional activity of the promoter ($s_1$ = off: the promoter cannot initiate transcription; $s_2$ := on: the promoter is prone to initiate transcription). In addition there are $n + 1$ "promoter regime" states, which we divide in two sets: $\{r_1, \ldots, r_n\}$ describe the 'low' 2-state promoter regime and the state $r_{n+1}$ describes the 'high' 2-state promoter regime. Transitions through the "promoter regime" states represent the regulatory processes that transmit regulatory information from the enhancer to the promoter. The promoter remains in the low regime until all the $n$ regulatory processes have been completed, and only at that point it can transition into the high regime (see Figure 1 in this document). The low and high regimes differ in their on, off and initiation rates . Finally, the number of RNA molecules per cell can be any integer $m \geq 0$.

We assume that the enhancer switches between its close and far states independently of the promoter and the promoter regime state. Transitions among the promoter regime states are reversible, however a forward transition is only possible when the enhancer is in the close state. Transition between the on and off state of the promoter are reversible and their rates depend on the promoter regime. Transcription can only be initiated from the promoter state $s_2$ (i.e on state, either in the low or high regime) and the initiation rate depends on the promoter regime. Synthesis and degradation of RNAs is regarded as a Poisson process (i.e. we describe RNA initiation, elongation, nuclear export of RNA as a single kinetic step). Note that "$n+1$ promoter regime states" actually means that there are $n$ intermediate regulatory steps between the low regime and the high regime. The kinetic reactions are as follows.

$$\begin{array}{rcl}
e_1 & \underset{k_{close}}{\overset{k_{far}}{\rightleftharpoons}} & e_2 \\
e_1 + r_k & \underset{k_{back}}{\overset{k_{forward}}{\rightleftharpoons}} & e_1 + r_{k+1}, \text{ for } \ k = 1, \ldots, n \\
e_2 + r_k & \underset{k_{back}}{\leftarrow} & e_2 + r_{k+1}, \text{ for } \ k = 1, \ldots, n \\
s_1 + r_k & \underset{k_{off}^{lo}}{\overset{k_{on}^{lo}}{\rightleftharpoons}} & s_2 + r_k, \text{ for } \ k = 1, \ldots, n \\
s_1 + r_{n+1} & \underset{k_{off}^{hi}}{\overset{k_{on}^{hi}}{\rightleftharpoons}} & s_2 + r_{n+1} \\
m + s_2 + r_k & \underset{\mu^{lo}}{\rightarrow} & m + 1 + s_2 + r_k, \text{ for } \ k = 1, \ldots, n \\
m + s_2 + r_{n+1} & \underset{\mu^{hi}}{\rightarrow} & m + 1 + s_2 + r_{n+1} \\
m & \underset{\delta}{\rightarrow} & m - 1
\end{array}$$

The chemical master equation is given by

$$\frac{dp}{dt}(e_i, s_j, r_k, m) = (m+1)\delta p(e_i, s_j, r_k, m+1) + \mu_{(i,j,k)} p(e_i, s_j, r_k, m-1) \quad (1)$$

$$+ \sum_{(\bar{i},\bar{j},\bar{k})} k_{(\bar{i},\bar{j},\bar{k};i,j,k)} p(e_i, s_j, r_k, m) \quad (2)$$

$$-(m\delta + \mu_{(i,j,k)} + \sum_{(\bar{i},\bar{j},\bar{k})} k_{(i,j,k;\bar{i},\bar{j},\bar{k})}) p(e_i, s_j, r_k, m) \quad (3)$$

where $\mu_{(i,j,k)}$ is the transcription rate given the enhancer state, promoter state and the promoter regime state, and $k_{(i,j,k;\bar{i},\bar{j},)}$ is the transition rate for the enhancer and promoter to go from the states $e_i, s_j, r_k$ to the states $e_{\bar{i}}, s_{\bar{j}}, r_{\bar{k}}$. All rates are expressed in terms of the model parameters and are defined in Table 1.

This model can be rephrased in the general framework of multi-state promoter models (Sánchez and Kondev [2008]) if we interpret the triplet $(e, s, r)$ as a "hyper" promoter state. There are $4(n+1)$ hyper promoter states which can be described either by a triplet $(i, j, k)$ where $e_i$ defines the enhancer state, $s_j$ defines the promoter state and $r_k$ defines the promoter regime state, or it can be described by an integer $\varphi \in \{1, 2, \ldots, 4(n+1)\}$. The two descriptors are connected by the so-called linear indexing bijection

$$(i, j, k) \to \varphi = f(i, j, k) := (i-1)2(n+1) + (j-1)(n+1) + k. \quad (4)$$

Using this new notation, The chemical master equation can be rewritten as

$$\begin{aligned}
\frac{dp}{dt}(\varphi, m) &= (m+1)\delta p(\varphi, m+1) + \mu_\varphi p(\varphi, m-1) \\
&\quad + \sum_{\vartheta} k_{(\vartheta;\varphi)} p(\varphi, m) \\
&\quad -(m\delta + \mu_\varphi + \sum_{\vartheta} k_{(\varphi;\vartheta)}) p(\varphi, m)
\end{aligned}$$

#### 2 Mean and variance of the number of RNA

Following Sánchez and Kondev [2008], we define the probability vector $\mathbf{p}(m) = [p(1,m), \ldots, p(4(n+1), m)]$ and rewrite the chemical master equation in matrix form

$$\frac{d\mathbf{p}}{dt}(m) = (\mathbf{K} - \mathbf{T} - m\mathbf{\Delta})\mathbf{p}(m) + (m+1)\mathbf{\Delta}\mathbf{p}(m+1) + \mathbf{T}\mathbf{p}(m-1) \tag{5}$$

where the matrix $\mathbf{K}$ has elements $K_{\varphi\vartheta} = k_{(f^{-1}(\varphi);f^{-1}(\vartheta))}$ if $\varphi \neq \vartheta$ and $K_{\varphi\varphi} = -\sum_{\vartheta\neq\varphi} K_{\varphi\vartheta}$; the matrix $\mathbf{T}$ is diagonal with diagonal elements $T_{\varphi\varphi} = \mu_{f^{-1}(\varphi)}$; and the matrix $\mathbf{\Delta}$ is also diagonal with diagonal elements $\mathbf{\Delta}_{\varphi\varphi} = \delta$. The mean and the variance of the number of RNAs per cell is given by the following formulas (Sánchez and Kondev [2008])

$$\langle \text{RNA} \rangle = \frac{\boldsymbol{\mu}\boldsymbol{m}^{(0)}}{\delta} \tag{6}$$

$$\text{Var(RNA)} = \frac{\boldsymbol{\mu}\boldsymbol{m}^{(0)}}{\delta} + \frac{\boldsymbol{\mu}\boldsymbol{m}^{(1)}}{\delta} - \left[\frac{\boldsymbol{\mu}\boldsymbol{m}^{(0)}}{\delta}\right]^2 \tag{7}$$

where we have defined the vector $\boldsymbol{\mu} = (\mu_{f^{-1}(1)}, \ldots, \mu_{f^{-1}(4(n+1))})$ and $\mathbf{m}^{(j)} = \sum_m m^j p(m)$ the $j$th moment of the number of RNAs per cell. The vectors $\boldsymbol{m}^{(0)}$ is the solution (subject to the normalisation $\sum_\varphi m_\varphi^{(0)} = 1$) of the equation

$$\boldsymbol{K}\mathbf{m}^{(0)} = 0. \tag{8}$$

(Sánchez and Kondev [2008]). The vectors $\boldsymbol{m}^{(1)}$ is the solution of the equation

$$(\mathbf{K} - \mathbf{\Delta})\mathbf{m}^{(1)} + \boldsymbol{T}\mathbf{m}^{(0)} = 0. \tag{9}$$

(see equation 4 in Sánchez and Kondev [2008]).

#### 3 Mean and variance of the number of RNAs as a function of the contact probability

In the enhancer-promoter model, the *contact probability* is the steady-state probability that the enhancer is in the close state. It can be directly calculated from the close rate and the far rate,

$$p_c = \frac{k_{close}}{k_{close} + k_{far}}. \tag{10}$$

From equation (10), we can express $k_{close}$ in terms of $k_{far}$ and $p_c$,

$$k_{close} = \frac{k_{far} p_c}{1 - p_c}. \tag{11}$$

We substitute equation (11) for $k_{close}$ in equation (5) and obtain the mean number of RNA molecules as a function of the rate parameters $k_{far}$, $k_{back}$, $k_{forward}$, $k_{on}^{lo}$, $k_{off}^{lo}$, $k_{on}^{hi}$, $k_{off}^{hi}$, $\mu^{lo}$, $\mu^{hi}$, the number of intermediate regulatory steps $n$, and the contact probability $p_c$.

### 4 Burst size and burst frequency

In our model, a transcriptional burst is defined as a "stay" of the promoter in the *on* state. Hence, the mean duration of a burst is the average time that the promoter stays in the *on* state, $\langle T_{on} \rangle$. The burst size is defined as the mean number of mRNA produced during an average burst, i.e.

$$b_s = \langle T_{on} \rangle (\mu^{hi} p_{on|high} + \mu^{low} p_{on|low}) \tag{12}$$

where $p_{on|high}$ and $p_{on|low}$ the steady-state probabilities that the promoter is in the *on* state given that it is in the low regime or in the high regime. The burst frequency is defined as the number of bursts per mRNA life time, i.e.

$$b_f = \frac{p_{on}}{\langle T_{on} \rangle \delta} \tag{13}$$

where $p_{on}$ is the steady-state probability that the promoter is in the *on* state.

The average duration of a stay in the *on* state can be directly calculated from the transition rate matrix $\mathbf{K}$ using the theory of absorbing Markov chain (Iosifescu [1980]).

### 5 Correlation between enhancer-promoter interactions and promoter activity

The correlation between enhancer-promoter interactions and promoter activity is the steady-state correlation between the enhancer state *close* and the promoter state *on*, i.e.

$$Corr(close, on) = \frac{p_{close,on} - p_{on} p_{close}}{\sqrt{p_{on}(1 - p_{on}) p_{close}(1 - p_{close})}} \tag{14}$$

where $p_{close,on}$ is the steady-state probability that the enhancer is in the *close*, state and that the promoter is in the *on*, state, $p_{close}$ is the steady-state probability that the enhancer is in the *close*, state, and $p_{on}$ is the steady-state probability that the promoter is in the *on*, state. Those three probabilities are calculated from the steady-state distribution of the hyper promoter states, $\mathbf{m}^{(0)}$, by summing its elements that correspond to the hyper promoter states $(close,on)$, *close*, and *on*, respectively.

### 6 Distribution of the number of RNAs per cell

The joined steady-state distribution of the enhancer state, promoter state, promoter regime state and number of RNA per cell, $\tilde{\mathbf{p}}(\cdot)$, is the equilibrium of equation (5), i.e. $\tilde{\mathbf{p}}(\cdot)$ solves

$$\frac{d\mathbf{p}}{dt}(m) = 0, \text{ for all } m. \tag{15}$$

The marginal steady-state distribution of the number of RNA per cell is given by

$$p^*(m) = \sum_{\varphi=1}^{4(n+1)} \tilde{p}_\varphi(m). \tag{16}$$

We calculated the joined steady-state probability distribution of the enhancer-promoter model by using the finite state projection method (Munsky and Khammash [2006]). This method consists of truncating the infinite state space of the system into a finite subset of states in order to reduce the infinite-dimensional system of ODEs into a finite system. For the enhancer-promoter model, this truncation consists in fixing a maximal number of RNAs per cell which we set to $120\%$ of the maximal number observed in the FISH experiment. The code for these calculations was written in Matlab (version 2019b) and are available on Github (`https://github.com/gregroth/Zuin_Roth_2021`).

### 7 Qualitative study of the transcriptional response to changes in contact probabilities

In this Section, we investigate the effect of the model parameters on the shape of the transcriptional response, i.e. the curve describing how enhancer-promoter contact probabilities are translated into mean number of RNAs. We will show that there are only three possible scenarios that we call *sub-linear*, *sigmoidal-like*, and *super-linear*. We define the curve to be sub-linear if its first derivative is positive for all contact probabilities and its second derivative is negative at contact probabilities $0$ and $1$. The curve is sigmoidal-like if its first derivative is positive and its second derivative is negative a contact probability $0$ and positive at contact probability $1$. Finally, the curve is super-linear if its first derivative is positive for all contact probabilities and its second derivative is positive at contact probabilities $0$ and $1$. Figure 2B illustrates the types of curve.

The main effect of the contact probability on the mean number of RNAs is due to the fact that enhancer-promoter contacts allow for a transient shift to the high promoter regime, which has a larger apparent transcription rate compared to the low promoter regime. Therefore, the relationship between contact probabilities and mean number of RNA (i.e. $\langle \text{RNA} \rangle (p_c)$) should be qualitatively similar to the relationship between contact probability and the probability that the promoter is in the high regime mode, i.e. $p_{hi}(p_c) = \mathbb{P}(r = n + 1)(p_c)$.

The high regime probability $p_{hi}(p_c)$ depends only on $4$ parameters (i.e. $k_{far}, k_{back}, k_{forward}$, and $n$). We can explore analytically the effect of each parameter in the shape of the transcriptional response. The high regime mode probability is directly calculated from the steady state probability vector $\mathbf{m}^{(0)}$ by summing its elements that correspond to the high regime mode (i.e. $(i, j, n + 1)$ for $i = 1, 2$ and $j = 1, 2$):

$$p_{hi} = \mathbf{e}^{hi}\mathbf{m}^{(0)} \tag{17}$$

where $\mathbf{e}^{hi}$ is the vector of length $4(n + 1)$ whose elements $e^{hi}_{f(i,j,k)} = 1$ are $1$ if $k = n + 1$ and $0$ elsewhere. The expression of $p_{hi}$ in terms of the model parameters is derived by solving the linear equation (8).

**Without intermediate regulatory steps ($n = 1$).** We first focus on a version of the model where the transition between the promoter low and high regimes occurs in a single step. $n = 1$. We show that in this case, the curve $p_{hi}(p_c)$ is always sub-linear.

The exact formula for $p_{hi}(p_c)$ is

$$p_{hi}(p_c) = \frac{p_c k_{forward}(k_{back} + k_{far} - p_c k_{back})}{p_c(k_{far}k_{forward} - k_{back}k_{forward} - k^2_{back}) + k_{back}k_{far} + k_{back}k_{forward} + k^2_{back}} \tag{18}$$

(we used the *symbolic* toolbox in *Matlab* to solve equation (8)). By defining the two dimensionless parameters $z = \frac{k_{far}}{k_{back}}$ and $u = \frac{k_{forward}}{k_{back}}$, we can rewrite equation (18) as

$$p_{hi}(p_c) = \frac{-up_c^2 + u(1+z)p_c}{p_c(zu - u - 1) + z + u + 1}. \tag{19}$$

In order to study the shape of $p_{hi}(p_c)$, we calculate its first and second derivatives

$$\frac{\partial p_{hi}}{\partial p_c}(p_c) = \frac{u(z^2 + 2(1-p_c)z + (1-p_c^2)uz + (1-p_c)^2 u + (1-p_c)^2)}{(z + u(1-p_c) + p_c zu + 1 - p_c)^2} \tag{20}$$

$$\frac{\partial^2 p_{hi}}{\partial p_c^2}(p_c) = -\frac{2z^2u^2(z+u+1)}{(z + u(1-p_c) + p_c zu + 1 - p_c)^3} \tag{21}$$

The first derivative $\frac{\partial p_{hi}}{\partial p_c}(p_c)$ is always positive and the second derivative $\frac{\partial^2 p_{hi}}{\partial p_c^2}(p_c)$ is always negative. Hence, $p_{hi}$ is a sub-linear function of the contact probability for any values of the parameters $u$ and $z$.

We then analyse how the parameters $u$ and $z$ affect the "strength" of the sub-linearity, which is related to how the probability of being in the high regime increases at low contact probability (i.e. $\frac{\partial p_{hi}}{\partial p_c}(0)$) and at high contact probability (i.e. $\frac{\partial p_{hi}}{\partial p_c}(1)$). Sub-linearity is "strong" when the probability of being in the high regime sharply increases at low contact probability (i.e. $\frac{\partial p_{hi}}{\partial p_c}(0)$ large) and plateaus at high contact probability (i.e. $\frac{\partial p_{hi}}{\partial p_c}(1)$ small). The derivatives at contact probabilities 0 and 1 are given by

$$\frac{\partial p_{hi}}{\partial p_c}(0) = \frac{u(1+z)}{(1+z+u)} \tag{22}$$

and

$$\frac{\partial p_{hi}}{\partial p_c}(1) = \frac{u}{(1+u)^2} \tag{23}$$

Thus the degree of sub-linearity increases when $u$ and/or $z$ increase, which means an increase in the ratio $\frac{k_{forward}}{k_{back}}$ and/or in the ratio $\frac{k_{far}}{k_{back}}$. Thus when memory is long, i.e. the promoter remains in the high-regime mode much longer than the average duration of an interaction ($\frac{k_{far}}{k_{back}} >> 1$), and the intermediate regulatory steps are fast ($\frac{k_{forward}}{k_{back}} >> 1$), then transcription levels are highly sensitive to changes in contact probabilities when contact probabilities are low, and conversely poorly sensitive when contact probabilities are high.

**With one intermediate regulatory step ($n = 2$).** When the number of intermediate regulatory steps is larger than 1, the formulae become more complicated. In order to understand the effect of the intermediate regulatory steps on the qualitative shape of the function $p_{hi}(p_c)$, we first restrict ourselves to the case $n = 2$. We show that for any value of the parameter $z$, there exist $u_1^* < u_0^*$ such that $p_{hi}(p_c)$ is sub-linear for $u \leq u_1^*$, sigmoidal-like for $u_1^* < u < u_0^*$ and super-linear for $u \geq u_0^*$ (see Figure 2A-B for examples).

For any value of the parameters, the derivative of $p_{hi}(p_c)$ is positive as the only way to reach the high regime is to move forward in the regulatory steps which is only possible when the enhancer and the promoter are in contact. Therefore, we only need to calculate the curvature at contact probability 0 and 1 which yields

$$\frac{\partial^2 p_{hi}}{\partial p_c^2}(0) = \frac{2u^2z^2(-u^2 + (z+2)u + (1+z)^2)}{(z^2 + 2uz + 2z + u^2 + u + 1)^3} \tag{24}$$

$$\frac{\partial^2 p_{hi}}{\partial p_c^2}(1) = -\frac{2u^2(u^4 + (2+z)u^3 + (1+3z)u^2 - (1+z))}{z(u^2+u+1)^3} \tag{25}$$

We now demonstrate our claim, i.e. the existence of $u_0^*$ and $u_1^*$ such that $\frac{\partial^2 p_{hi}}{\partial p_c^2}(0) < 0$ and $\frac{\partial^2 p_{hi}}{\partial p_c^2}(1) < 0$ for $u \leq u_1^*$, $\frac{\partial^2 p_{hi}}{\partial p_c^2}(0) > 0$ and $\frac{\partial^2 p_{hi}}{\partial p_c^2}(1) < 0$ for $u_1^* < u < u_0^*$ and $\frac{\partial^2 p_{hi}}{\partial p_c^2}(0) > 0$ and $\frac{\partial^2 p_{hi}}{\partial p_c^2}(1) > 0$for $u \geq u_0^*$. The sign of $\frac{\partial^2 p_{hi}}{\partial p_c^2}(0)$ depends only on the second factor in the numerator of equation (24) which has a single positive root,

$$u_0^* = 1 + \frac{z}{2} + \sqrt{(z+2)(z+1)}. \tag{26}$$

The sign of $\frac{\partial^2 p_{hi}}{\partial p_c^2}(1)$ depends only on the second factor in the numerator of equation (25) which has a single positive root, $u_1^*$ (in theory its expression can be calculated but it is not needed here). Next we show that $u_1^* < u_0^*$. Since the the second factor in the numerator of equation (25) is an increasing function in $u$ and that its sign is negative when $u = 0$, we only need to show that its sign is positive for $u = u_0^*$. This is the case because $2 < 1 + \frac{z}{2} + \sqrt{(z+1)(z+1)} < u_0^*$ and the second factor in the numerator of equation (25) evaluated in $u = 2$ is clearly positive for any value of $z$. The signs of $\frac{\partial^2 p_{hi}}{\partial p_c^2}(0)$ and $\frac{\partial^2 p_{hi}}{\partial p_c^2}(1)$ as function of $u$ and $z$ are shown in Figure 2A.

**With more intermediate steps ($n > 2$).** Finally, we conjecture that this trichotomy (i.e. sub-linear, sigmioal-like, super-linear) is still valid for small $n > 2$ but the sub-linear parameter region tends to disappear for large $n$. This is motivated by numerical calculations of the second derivatives $\frac{\partial^2 p_{hi}}{\partial p_c^2}(0)$ and $\frac{\partial^2 p_{hi}}{\partial p_c^2}(1)$ for a large set of values for the parameters $u$ and $z$ and for different values of $n$ (see Figure 3A-B for few examples). Thus, we investigate numerically the effect of $n$ on the sensitivity of the curve at low contact probability which we measure as $\frac{\partial p_{hi}}{\partial p_c}(0)/p_{hi}(1)$. For that we approximate numerically the derivative at contact probability 0 and vary the number of regulatory steps $n$ and the dimensionless parameter $u$ and $z$. Figure 3C shows that the sensitivity increases with an increase in the number of regulatory steps.

### 8 Model fitting

The enhancer-promoter communication model described above was fitted simultaneously to the mean eGFP levels measured in individual cell lines, to the distribution of RNA numbers measured by smRNA FISH in a control line where the SCR is absent ($C^l$), and in a cell line where the full-length Sox2 control region (SCR) is adjacent to the promoter ($C^h$). This section describes in detail how this was done.

#### 8.1 RNA FISH data.

For the cell lines $C^k$ (i.e.$k = l, h$), we calculated the histogram, $\mathbf{h}^k$ of the RNA molecule counts obtained from the smRNA FISH experiment. The bin size $b_k$ and the number of bins $n_b^k$ were chosen using the function *histogram* in Matlab.

For each set of rates $\boldsymbol{\theta} = (k_{far}, k_{back}, k_{forward}, k_{on}^{lo}, k_{off}^{lo}, k_{on}^{hi}, k_{on}^{hi}, \mu^{lo}, \mu^{hi})$ and each number of regulatory steps $n$, we calculated the steady-state probability distributions predicted by the enhancer-promoter model with parameters $(n, \boldsymbol{\theta})$ and contact probabilities equal to 0 (for the cell line $C^l$) and 1 (for the cell line $C^h$). Next, we discretised the steady-state distributions in a histogram $\boldsymbol{\eta}^k(n, \boldsymbol{\theta})$ which is comparable with the histogram $\mathbf{h}^k$ obtained from the FISH data. We assume that the count in bin $i$ for cell line $k$ follows a binomial distribution of mean $\eta_i^k N_k$ and variance $\eta_i^k(1-\eta_i^k)N_k$, where $N_k$ is the total number of counts. We approximate the binomial distribution by a normal distribution. Hence, the likelihood of observing the histogram $\mathbf{h}^k$ given the parameters $(n, \boldsymbol{\theta})$ is

$$L_k(n,\boldsymbol{\theta}) = \prod_{i=1}^{n_b^k} \frac{1}{\sqrt{2\pi N_k \eta_i^k(1-\eta_i^k)}} e^{-\frac{(h_i^k-\eta_i^k(\boldsymbol{\theta}))^2}{2\eta_i^k(1-\eta_i^k)N_k}} \tag{27}$$

and the log likelihood is

$$LL_k(n,\boldsymbol{\theta}) = \sum_{i=1}^{n_b^k} \left[ -\frac{(h_i^k-\eta_i^k(\boldsymbol{\theta}))^2}{2\eta_i^k(1-\eta_i^k)N_k} + \frac{1}{2}\log(2\pi N_k \eta_i^k(1-\eta_i^k)) \right] \tag{28}$$

#### 8.2 Mean eGFP levels.

The data consist of the inferred mean number of RNA molecule per cell in each of the $N$ individual eGFP+ cell lines (obtained via the calibration with sRNA FISH (Suppl. Fig. 1H)) and the associated genomic distance from the promoter to the SCR. The data were then averaged in bins of length 20 kb, yielding a vector g whose elements are the binned mean number of RNA per cell and a vector $\boldsymbol{d}$ of genomic distances. The genomic distances were transformed in contact probabilities using the Capture Hi-C data (6.4-kb resolution; see Figure 2A), yielding a vector $\boldsymbol{\pi}$ of contact probabilities associated to the vector $\boldsymbol{g}$.

For each set of rates $\boldsymbol{\theta} = (k_{far}, k_{back}, k_{forward}, k_{on}^{lo}, k_{off}^{lo}, k_{on}^{hi}, k_{on}^{hi}, \mu^{lo}, \mu^{hi})$, each number of regulatory steps $n$, and each cell line $i$, we calculate the mean number of RNA per cell, $\gamma_i(n, \boldsymbol{\theta}, \pi_i)$, predicted by the enhancer-promoter model with parameter $(n, \boldsymbol{\theta})$ and contact probability $\pi_i$ (using equation (6)). Assuming that the deviations from the model are normally distributed with mean 0 and variance $\sigma^2$, the log likelihood function is given by

$$LL_{mean}(n,\boldsymbol{\theta}) = -\sum_{i=1}^{N} \frac{(g_i - \gamma_i(\boldsymbol{\theta}, \pi_i))^2}{2\sigma^2} + N\frac{1}{2}\log(2\pi\sigma) \tag{29}$$

where $N$ is the number of cell lines.

#### 8.3 Model fitting for the full-length SCR data set

When the contact probability is 0, the enhancer-promoter model reduces to a two-state model with parameters $\boldsymbol{\theta}_l = (k_{on}^{lo}, k_{off}^{lo}, \mu^{lo})$. Therefore, we first fit a two-state model to the RNA FISH distribution of the control line where the SCR is absent ($C^l$). The best fit parameters of the low regime $\boldsymbol{\theta}_l^*$ maximises the log likelihood function $LL_l(n, \boldsymbol{\theta})$ where all parameters other than the ones in $\boldsymbol{\theta}_l$ are set to a fixed value.

Next we fit the enhancer-promoter model simultaneously to both the binned mean eGFP levels measured in individual cell lines and the RNA FISH distribution of the cell line where the SCR is adjacent to the promoter ($C^h$), keeping the low regime parameters fixed to their best fit values, i.e. $\boldsymbol{\theta}_l^*$. The best fit parameter $(n^*, \boldsymbol{\theta}^*)$ maximises the total log likelihood function

$$LL_{tot} := LL_{mean}(n,\boldsymbol{\theta}) + LL_h(n,\boldsymbol{\theta}). \tag{30}$$

Since the number of regulatory steps $n$ is an integer parameter, we maximise the total log likelihood function $LL_{tot}$ for each value of $n$ separately. For the parameter $n$, we set a lower bound of 1 and an upper bound of 10. For parameters $k_{far}, k_{back}, k_{forward}, k_{on}^{lo}, k_{off}^{lo}, k_{on}^{hi}, k_{off}^{hi}$, we set a lower bound of 0 and an upper bound of 100. Finally for the parameters $\mu^{lo}, \mu^{hi}$, we set a lower bound of 0 and an upper bound of 3000. We ensured that the best fit parameters found were not at the boundaries. For all the maximisations we use a global search approach. Specifically, we use the Matlab function *MultiStart* in the *Global Optimization* toolbox. All codes were written in Matlab (version 2019b) and are available at `https://github.com/gregroth/Zuin_Roth_2021`.

**Best fit parameter sensitivity.** For each best-fit parameter value, we numerically calculated its minimal perturbation required for a 0.02 increase in the $R^2$ of the model prediction. Figure 4 shows that the best fit is mostly sensitive to the rates driving the regulatory steps and to the high regime parameter rates. The best fit is weakly sensitive to the number of regulatory steps ($\Delta R^2 < 0.02$ for $4 \leq n \leq 11$).

#### 8.4 Model fitting for the truncated SCR data set

We fit the enhancer-promoter model to the binned mean number of RNA molecule inferred from the eGFP+ cell lines with the truncated version of the SCR. We successively maximise the log likelihood function $LL_{mean}$ under the constrain of having different combinations of free parameters – the other ones being fixed to the best fit values obtained for the full-length SCR data set. For each combination we calculate the $R^2$ of the model prediction. Finally, we rank all the combinations in terms of their $R^2$. Any combination of the parameters describing the high regime (i.e. $k_{on}^{hi}, k_{off}^{hi}, \mu^{hi}$) yields a $R^2 > .94$. Any combination of the parameters describing the intermediate steps processes (i.e. $k_{back}, k_{forward}$) yields a $R^2 < 0.22$. Finally, when $k_{on}^{hi}, k_{off}^{hi}, \mu^{hi}$ and $k_{back}, k_{forward}$ are all free parameters, $R^2$ is equal to $0.97$.

### References

M Iosifescu. *Finite Markov Processes and Their Applications*. Wiley, New York, New York, 1980.

Brian Munsky and Mustafa Khammash. The finite state projection algorithm for the solution of the chemical master equation. *The Journal of Chemical Physics*, 124(4):044104, 2006. doi: 10.1063/1.2145882.

Álvaro Sánchez and Jané Kondev. Transcriptional control of noise in gene expression. *Proceedings of the National Academy of Sciences*, 105(13):5081–5086, 2008. doi: 10.1073/pnas.0707904105.

| Rate | Expression | From | To |
|---|---|---|---|
| **Hyper state transition rates** | | | |
| $k_{(1,j,k;2,j,k)}$ for $j=1,2$, $k=1,\ldots,n+1$ | $k_{far}$ | close | far |
| $k_{(2,j,k;1,j,k)}$ for $j=1,2$, $k=1,\ldots,n+1$ | $k_{close}$ | far | close |
| $k_{(i,1,k;i,2,k)}$ for $k=1,\ldots,n$ | $k_{on}^{lo}$ | off/low | on/low |
| $k_{(i,2,k;i,1,k)}$ for $k=1,\ldots,n$ | $k_{off}^{lo}$ | on/low | off/low |
| $k_{(i,1,n+1;i,2,n+1)}$ for $i=1,2$ | $k_{on}^{hi}$ | off/high | on/high |
| $k_{(i2,n+1;i,1,n+1)}$ for $i=1,2$ | $k_{off}^{hi}$ | on/high | off/high |
| $k_{(1,j,k;1,j,k+1)}$ for $j=1,2$, $k=1,\ldots,n$ | $k_{forward}$ | close/$r_k$ | close/$r_{k+1}$ |
| $k_{(2,j,k;2,j,k+1)}$ for $j=1,2$, $k=1,\ldots,n$ | $0$ | far/$r_k$ | far/$r_{k+1}$ |
| $k_{(i,j,k;i,j,k-1)}$ for $j=1,2$, $k=2,\ldots,n+1$ | $k_{back}$ | $r_k$ | $r_{k-1}$ |
| **Initiation rates** | | | |
| $\mu_{i,2,k}$ for $i=1,2$, $k=1,\ldots,n$ | $\mu^{lo}$ | on/low regime | on/low+1 RNA |
| $\mu_{i,2,n+1}$ for $i=1,2$ | $\mu^{hi}$ | on/high regime | on/high +1 RNA |

**Table 1:** Transition rates used in equations (1) and (5). All the rates $\mu_{(i,j,k)}$ and $k_{(i,j,k;\bar{i},\bar{j},\bar{k})}$ that are not described in the table have value equal to $0$.

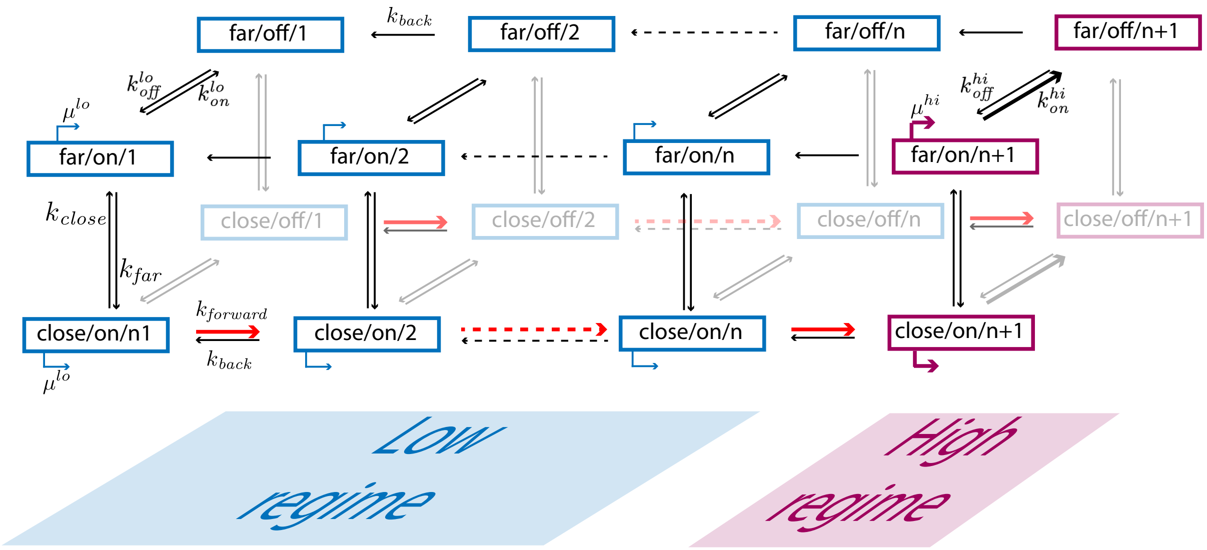


**Figure 1:** Scheme of the enhancer-promoter model. For simplicity every rate is indicated only once in similar reactions. Opacity differences are only intended to increase the clarity of the figure and do not relate to properties of the states themselves.

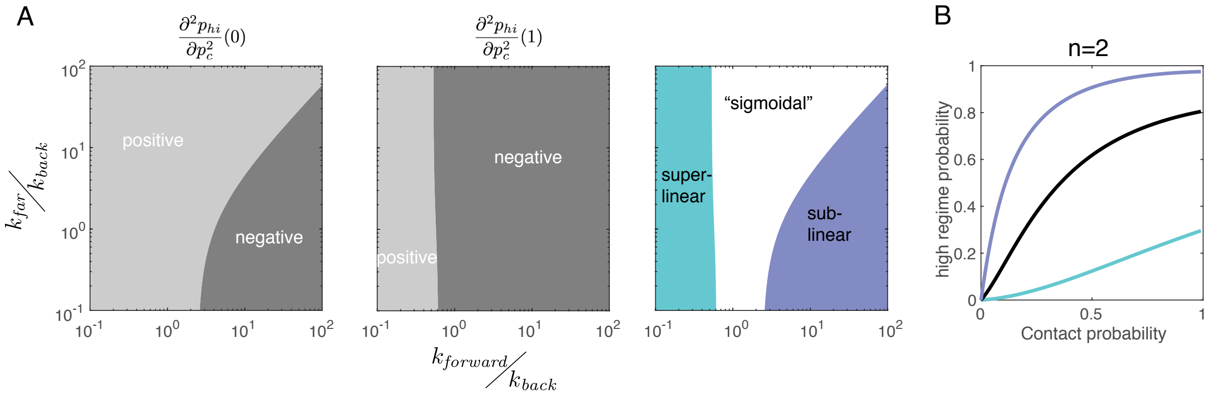


**Figure 2:** A. Sign of the second derivative of the probability of the promoter being in the high 2-state regime, plotted as a function of the dimensionless parameters $u = \frac{k_{forward}}{k_{back}}$ and $z = \frac{k_{far}}{k_{back}}$ when the contact probability is 0 (left panel) and 1 (centre panel), and the number of regulatory step is $n = 2$. The right panel shows the resulting trichotomy in the shape of the curve $p_{hi}(p_c)$. B. Representation of the three possible shapes of the curve $p_{hi}(p_c)$. Parameters: $n = 2, z = .1, u = .9$ (dark blue line), $u = 5$ (black line), $u = 40$ (light blue line).

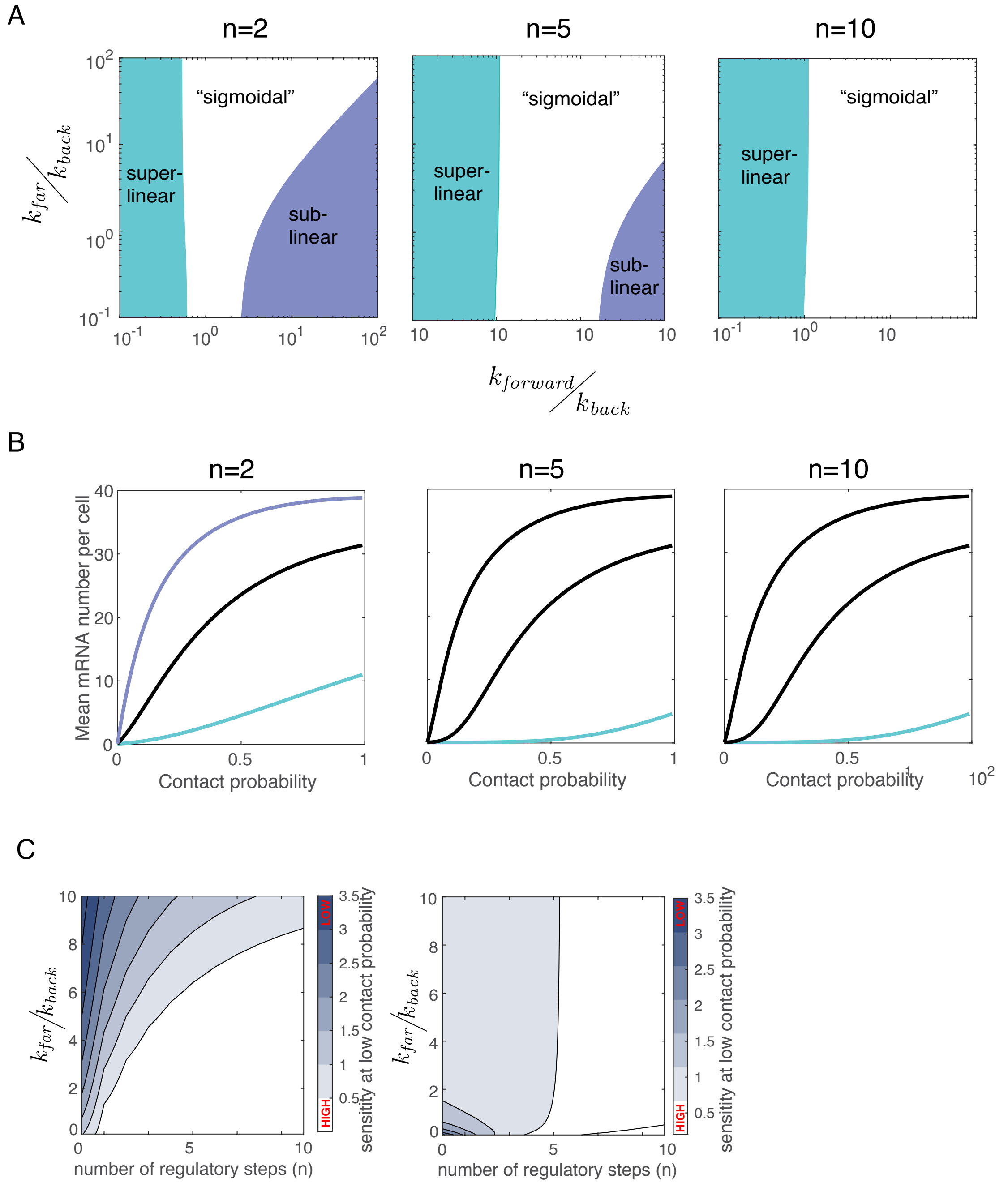


**Figure 3:** A. Types of the curve $p_{hi}(p_c)$ (i.e. sub-linear, sigmoidal-like, super-linear) as a function of the dimensionless parameters $u = \frac{k_{forward}}{k_{back}}$ and $z = \frac{k_{far}}{k_{back}}$ for different numbers of regulatory steps (left, $n = 2$; centre, $n = 5$; right, $n = 10$). B. Representation of mean number of RNA predicted by the enhancer-promoter model with parameters $k_{far}, k_{forward}$ corresponding to the parameters chosen in Figure 2B and the other parameters are $k_{back} = 1, k_{on}^{lo} = .1, k_{off}^{lo} = 1, k_{on}^{hi} = 4, k_{off}^{hi} = 2, \mu^{lo} = 1, \mu^{hi} = 60$. C. Sensitivity study of the curve $p_{hi}(p_c)$ at contact probability 0. The degree of sensitivity is calculated as $\frac{\partial p_{hi}}{\partial p_c}(0)/p_{hi}(1)$.

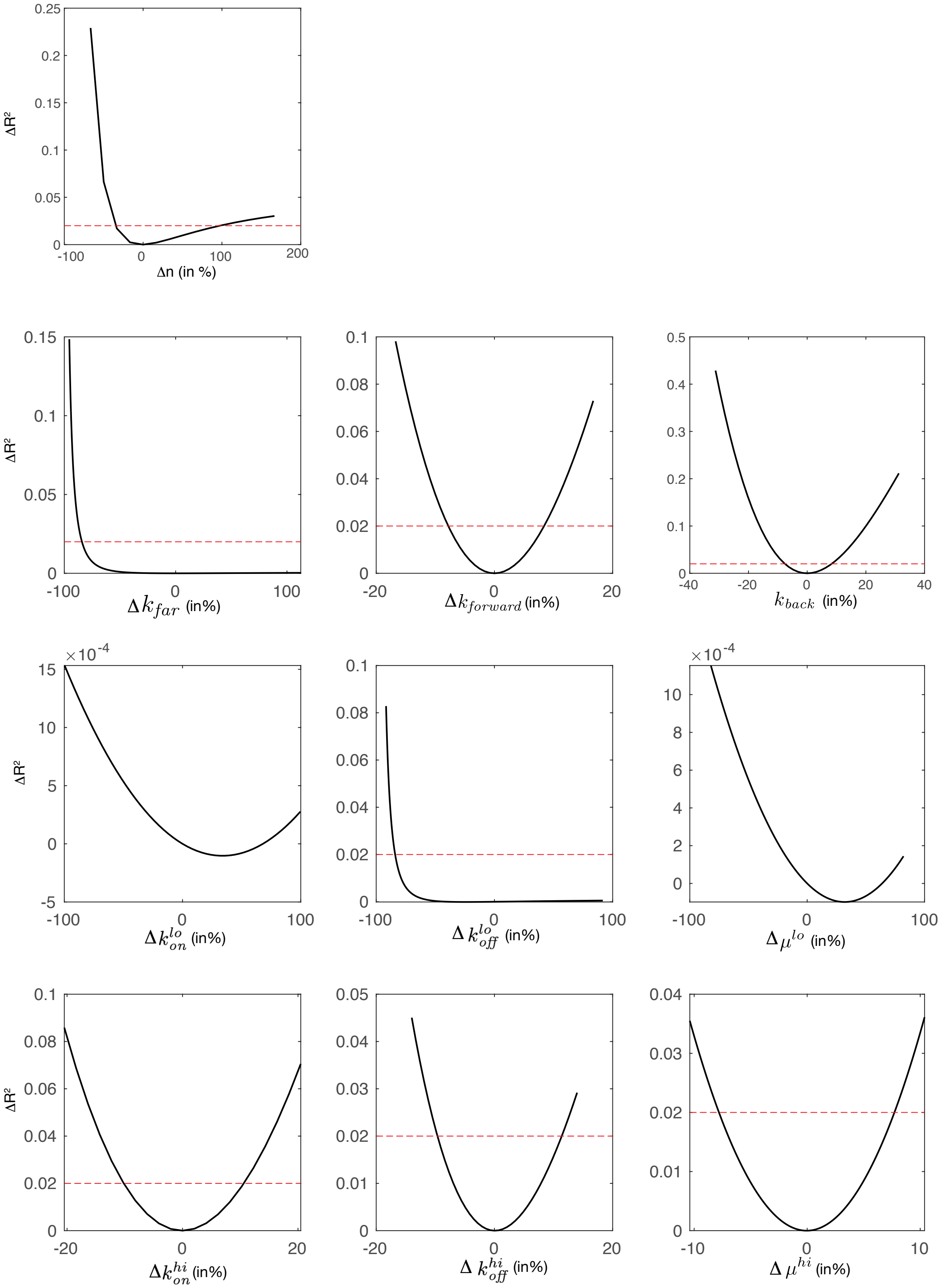


**Figure 4:** Parameter sensitivity. For each parameter, the difference in $R^2$ from best fit is plotted as a function of the difference in parameter value from best fit (in $\%$). The dashed red line represents a threshold at .02
